# Botanical Digitization: Application of MorphoLeaf in 2D Shape Visualization, Digital Morphometrics, and Species Delimitation, using Homologous Landmarks of Cucurbitaceae Leaves as a Model

**DOI:** 10.1101/2020.11.16.384230

**Authors:** Oluwatobi A. Oso, Adeniyi A. Jayeola

## Abstract

Morphometrics has been applied in several fields of science including botany. Plant leaves are been one of the most important organs in the identification of plants due to its high variability across different plant groups. The differences between and within plant species reflect variations in genotypes, development, evolution, and environment. While traditional morphometrics has contributed tremendously to reducing the problems that come with the identification of plants and delimitation of species based on morphology, technological advancements have led to the creation of deep learning digital solutions that made it easy to study leaves and detect more characters to complement already existing leaf datasets. In this study, we demonstrate the use of MorphoLeaf in generating morphometric dataset from 140 leaf specimens from seven Cucurbitaceae species via scanning of leaves, extracting landmarks, data extraction, landmarks data quantification, and reparametrization and normalization of leaf contours. PCA analysis revealed that blade area, blade perimeter, tooth area, tooth perimeter, height of (each position of the) tooth from tip, and the height of each (position of the) tooth from base are important and informative landmarks that contribute to the variation within the species studied. Our results demonstrate that MorphoLeaf can quantitatively track diversity in leaf specimens, and it can be applied to functionally integrate morphometrics and shape visualization in the digital identification of plants. The success of digital morphometrics in leaf outline analysis presents researchers with opportunities to apply and carry out more accurate image-based researches in diverse areas including, but not limited to, plant development, evolution, and phenotyping.

## INTRODUCTION

Over the years, many quantitative analytical tools have been used to analyse the diversity of plant structures, and morphometrics has become one of the most popular method in these analyses (Manacorda and Asurmendi, 2018). It deals with the study of variations that exist in shapes and biologically relevant structures, and it has been applied to multiple fields including plant systematics, plant development and evolution, zoology, geology, geography, and other fields that depend on comparison of structures, outlines, and contours(Hernández-Esquivel et al., 2020; Itgen et al., 2019; Pérez-Miranda et al., 2020; Terhune et al., 2020).

In plants, leaves are one of the most important features in plant identification (reference), and they present a worthy model to study diversity and pattern of evolution in plants via morphometry. The diversity of leaf forms is a result of multiple factors including genetic sequence, molecular pathways, developmental patterns, and the environment (Chitwood and Sinha, 2016; Dkhar and Pareek, 2014; Edwards et al., 2016; Ichihashi et al., 2014; Nicotra et al., 2011; Royer et al., 2009). Therefore, to understand the variation in leaf forms, it is important to do an in accurate analysis of the different leaf landmarks (Page et al., 2015; Soltis, 2017; Willis et al., 2017).

Depending on the dataset used, Morphometrics have been done using three methods including traditional morphometrics, outline-based morphometrics, and landmark-based morphometrics. Traditional morphometrics have focused mostly on basic measurements like length, breath, and angles, outline-based morphometry is the summary of the shape outlines without the landmarks, while landmark-based morphometry is the summary of the shape on the basis of its landmarks. Geometric morphometrics (GMM) however combines both landmark and outline analysis in studying diversity and shape variation within species (Cope et al., 2012; Punyasena and Smith, 2014). GMM allows for reconstruction of average leaf shape and a visualization of the morphospace in which each species leaf shape belongs (Alasadi et al., 2017; Borges et al., 2020; Fu et al., 2017; Klein and Svoboda, 2017; Lexer et al., 2009; Li et al., 2018)

Multiple programs have been used to examine the geometric morphometrics of leaves such as geomorph (Adams and Otárola-Castillo, 2013), tpsUtil and tpsDig2 (Rohlf, 2015), MorphoJ (Klingenberg, 2011), ImageJ (Abramoff et al., 2004), LEAFPROCESSOR (Backhaus et al., 2010) MorphoJ (Klingenberg, 2011), MorphoLeaf (Biot et al., 2016), and MASS (Chuanromanee et al., 2019), to analyze the landmark data. The present study uses MorphoLeaf (Biot et al., 2016) to integrate all basic steps of GMM analysis with an all-in-one method of preserving the leaf outline and integrity, and extracting all the vital details of the leaf details for multiscale analysis. It identifies biologically relevant and homologous landmarks along the leaf outlines with the aim of computing mean shapes. MorphoLeaf is available as a plug-in for Free-D software (Andrey and Maurin, 2005).

This work is an investigation into common exploratory and confirmatory techniques in landmark-based geometric morphometrics. The Curcubitaceae family presents itself as a great opportunity to for effective outline visualization in combination with principal component analysis and summarization of shape variables, due to the qualitative diversity reflected in its leaves. This led us to asking two questions (1) what heritable structural differences and similarities will be revealed from the foliar designs in the Curcubitaceae family vis-à-vis what are the evidences suggested about these homologous landmarks? And (2) do naturally homologous points in Curcubitaceae leaves allow trait diversity to be quantitatively tracked, and applied to functionally integrate systematics and digital identification of plants?

## MATERIALS AND METHODS

### Ethics Statement

No specific permits were required for the described field studies: a) no specific permissions were required for these locations/ activities; b) location are not privately-owned or protected; c) the field studies did not involve endangered or protected species.

### Sampling

Seven species of the Cucurbitaceae family were considered on the basis of common leaf shape and patterns, to investigate variations within and among individuals leaves, analyse the outline of the biological object or on biological landmarks, and compare the results of the Procrustes analysis to those of elliptical Fourier analysis.

They were collected in the wild so as to exclude the bias possibility that may set in with domestication. From simple to complex shapes, they include Lagenaria siceraria, Coccinia grandis, *Cucurbita pepo, Benincasa hispida, Trichosanthes cucumerina, Momordica charantia, and Citrullus colocynthis*.

Approximately 50 mature leaves were collected per species from different populations, but only twenty leaves per species (n=20) were scanned after careful selection so that each species contribute equal weight to sampling size. While there might be differences in sizes among species within the family, we focused on leaf shapes produced with identifiable homologous landmarks and noticeable differences in outlines.

#### Imaging

For scanning, an HP Laserjet Pro M1136 Mono Multi-function Laser Printer was used. A 20-cm metallic ruler was positioned at the side of each scanned sheet as a size marker. Leaves were placed directly on the scanner, adaxial face down, and scanned at a resolution of 300 dpi. Fresh leaves were used for the most part, however we used one-week old pressed specimen for *Coccinia grandis* and *Momordica charantia*

### Landmark Data Extraction

We followed the guidelines in the MorphoLeaf manual involving several steps including contour extraction, sinuses and peaks extractions, data extraction, data quantification, data normalization, and data representation (Biot et al., 2016).

Previous works used 17 (Chitwood et al., 2016a) and 21 (Chitwood et al., 2016b), but the number of biologically relevant landmarks we selected for this work was 17 with a few landmarks different from what has been used in previous works.

#### Extraction of the leaf contour (or outline)

Here, the watershed method employed by the MorphoLeaf application automatically removes biologically nonrelevant details along the contour, by retaining only the first elliptical Fourier descriptors that encode the contour in the frequency domain (Kuhl and Giardina, 1982). Due to the quality of the images, the number of descriptors was enough to automatically retain the fidelity of the contour after extraction. The leaf contour was automatically extracted using and manually corrected where necessary, particularly for leaves with unclear borders mostly due to deep sinuses. During the next step, two landmarks corresponding to the petiole were set manually, which allowed the automatic identification of the blade and the petiole. The leaf tip was then automatically determined as the point of the blade contour furthest away from the midpoint between petiole landmarks. This also defined the base-tip axis separating the blade in two half blades.

#### Identification of sinuses and tips of teeth

In the next step, we automatically identified the teeth, which are defined as portions of the blade contour between two sinuses – they are otherwise called lobes as in the Cucurbitaceae. Sinuses, which correspond to contour points with a high concave curvature, are identified in a two-step procedure. First, candidate intervals of the contour are determined as continuous domains where the curvature remains concave and above a user-defined threshold. Second, within each candidate interval, the point with the maximal curvature is selected as a sinus. After automatic detection of sinuses, errors were manually corrected to ensure that tooth limits are correctly positioned and avoid biases in subsequent analyses. After sinus identification, MorphoLeaf determined the position of the tooth tip between consecutive sinuses. The user can choose one of the two strategies available depending on tooth shape, however, we chose the maximum local curvature.

Although, identification of teeth hierarchy is crucial for proper characterizations when there are serrations on the leaves, we did not do it because the hierarchy of palmately lobed leaves with several levels of dissection cannot be established with MorphoLeaf. However, appropriate setting of the parameters was enough for detection of the sinuses on the main lobes and thus proper quantitative analyses and mean leaf shape reconstruction

#### Shape analysis (Elliptical Fourier Analysis)

Here we extracted the quantitative parameters of each leaf per species. The measures of the leaf contours, petiole/blade junctions, teeth sinuses, teeth peaks, and leaf apex landmarks had been validated. It was therefore processed to generate two files containing the quantifications of 17 landmark parameters including blade length, blade width (bb), blade width (is), blade area, blade perimeter, petiole width, upper teeth number, lower teeth number, total teeth number, tooth position, tooth width, tooth height (latitude), tooth length (median), tooth area, tooth perimeter, tooth position from the tip, tooth position from the base

#### Normalization

Here we reconstructed average leaf shapes using the second method in the guide. First, we resampled the leaves contour by using parameters of only two landmarks – leaf apex and petiole junction. We didn’t include the other primary sinuses and tips because it is optional (i.e. it doesn’t change the contour resamples), it and also caused difficulty in species with really high number of vertices. Reducing vertices thus meant that the shape outline might not be accurate. We then computed the moving average shapes of leaves. We define the Neighbor Rank as 20, corresponding to the number of leaves that contributed to each average shape and to produce a smoother overall form of normalized shape.

### Morphometric Analyses

After the extraction of homologous landmark and shape analysis was done, principal component analyses and correlation analyses was done on the dataset to compare the statistical differences between the variable and distribution of landmarks across the species. All statistical functions were performed in R (R Development Core Team, 2011) and visualized in the package ggplot2 (Wickham, 2016).

## RESULTS

### Landmark and extraction of leaf data

The primary basis for using MorphoLeaf is to automatically extract homologous landmark data for analysis. This included all the processes of landmarking, sinus and peak selection, landmark editing and correction, and data averaging and normalization. The 140 leaves analyzed in this study come from seven different species of Cucurbitaceae representing seven genera. While these shapes are mostly palmate with several primary and secondary teeth (Fig. 1a), it excludes secondary leaf types with leaflets. MorphoLeaf automatically extracted some landmarks of the leaves, however, editing and manual correction was done on few others because of the overlap in some of the teeth. The dataset subsequently extracted were then used for reparametrisation or generation of mean shapes after normalization. Secondarily, the reconstructed leaf shapes can be visualized using sviewer, a standalone version of Free-D’s 3D/2D rendering module for viewing curvature (“.cv”) file. The visualization of the mean shapes outlines after reparametrisation is captured in Fig. 1b.

**Fig. 1A:**
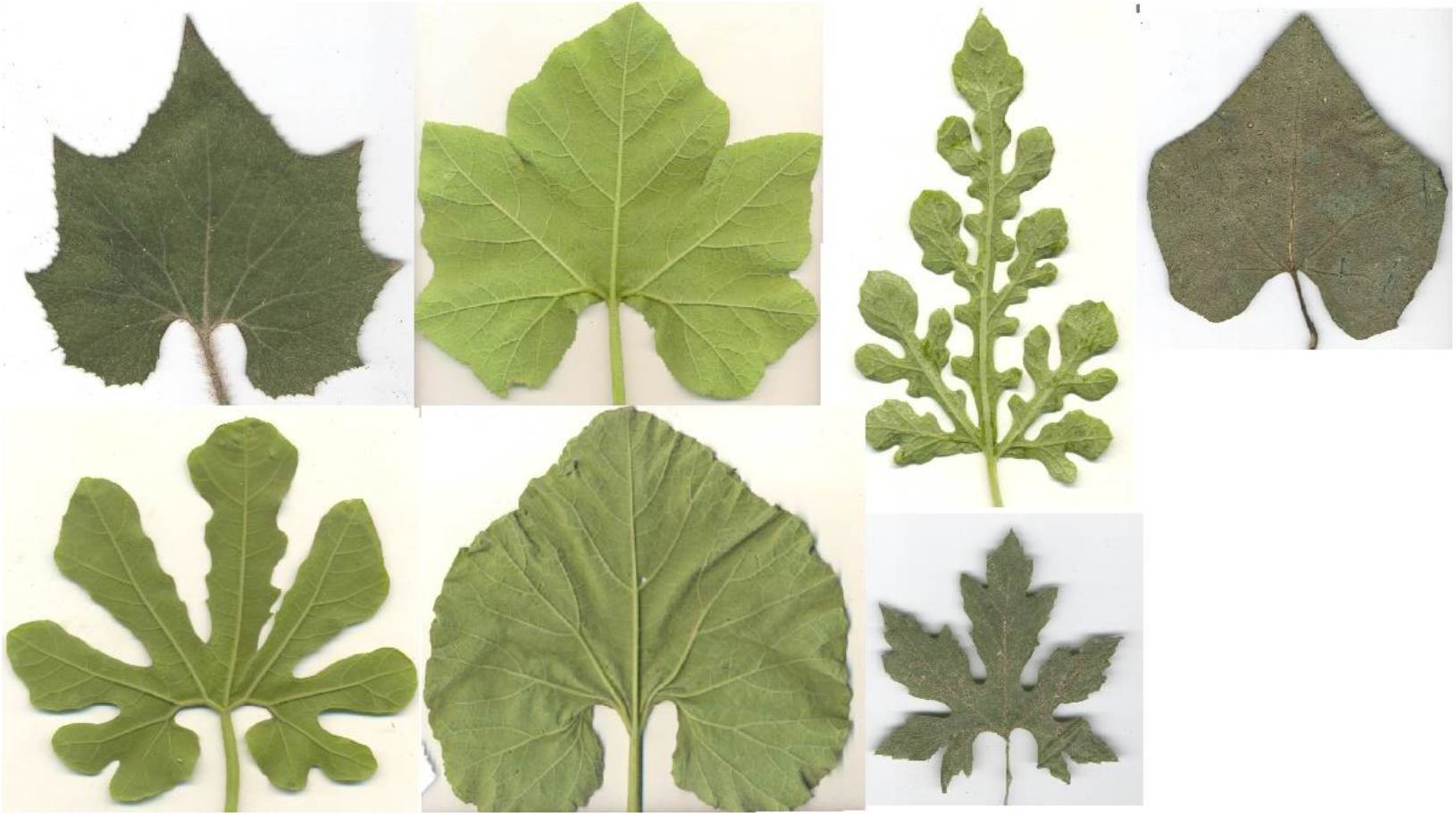
Seven species of Cucurbitaceae representing seven genera. Representative of theleaf specimen studied, showing the leaf outline and teeth after scanning, before processing and analysis. *Benincasa hispida, Cucurbita pepo, Citrullus colocynthis, Trichosanthes cucumerina, Lagenaria siceraria, Momordica charantia*, and *Coccinia grandis* (From top, left to right).

**Fig. 1B:**
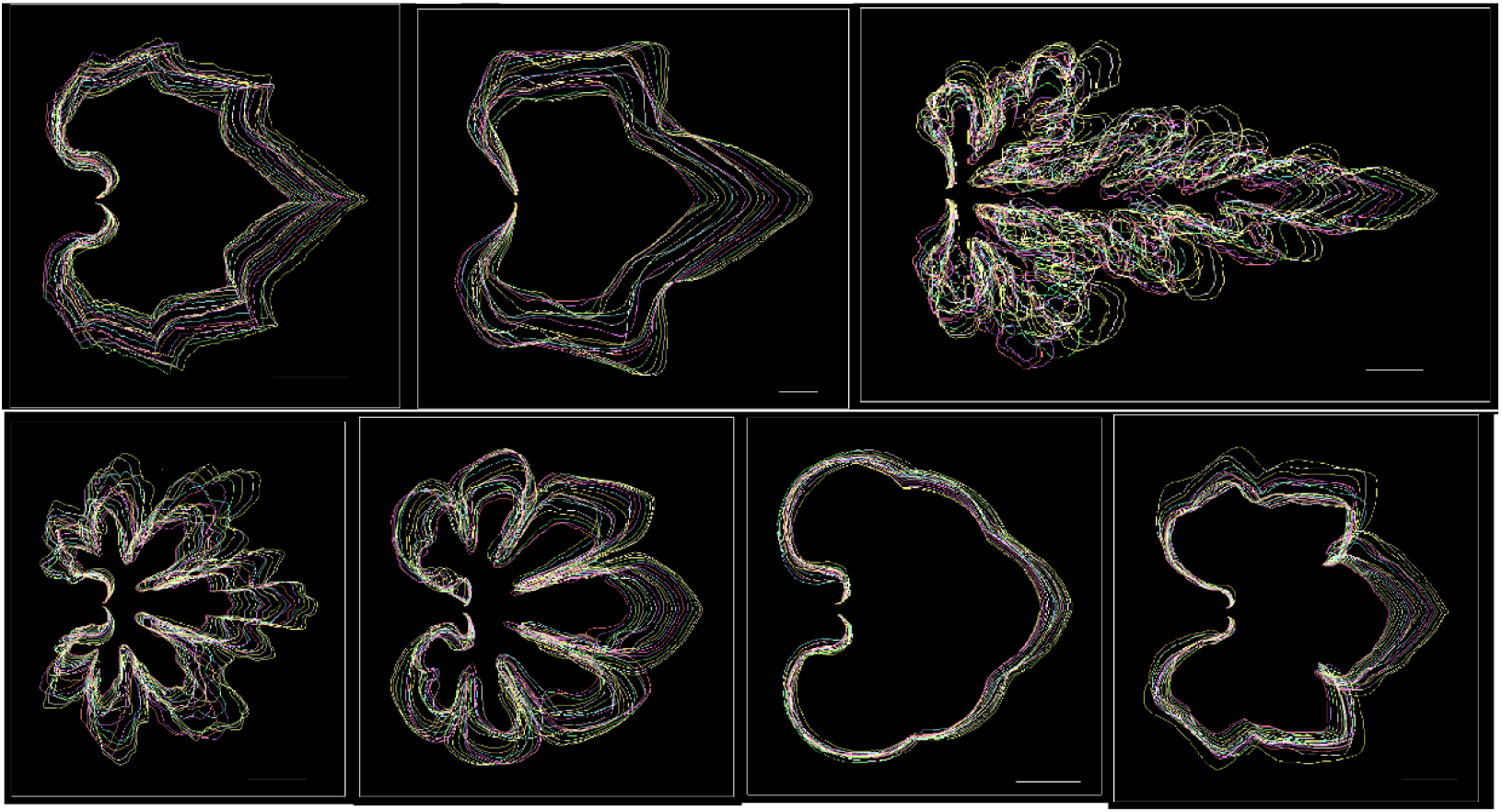
Mean shapes visualization after landmarking. After landmarking and reparametrization of biologically relevant points, the mean shapes are visualized to view the stack of each species’ contours. *Benincasa hispida, Coccinia grandis, Citrullus colocynthis, Momordica charantia, Trichosanthes cucumerina, Lagenaria siceraria*, and *Cucurbita pepo* (From top, left to right).

### Leaves Measures Analysis

PCA analysis was first carried out on the leaf dataset excluding the teeth dataset to see the quantitative differences in the width (BB and IS), length, area, and perimeter. The measures obtained from the outlines of these species also grouped them into seven. The coefficient matrix of the leaves’ dataset (Fig. 2a) showed a strong positive correlation between these five variables across all seven species, but the distributions of each variables are statistically different between the species.

**Fig. 2A:**
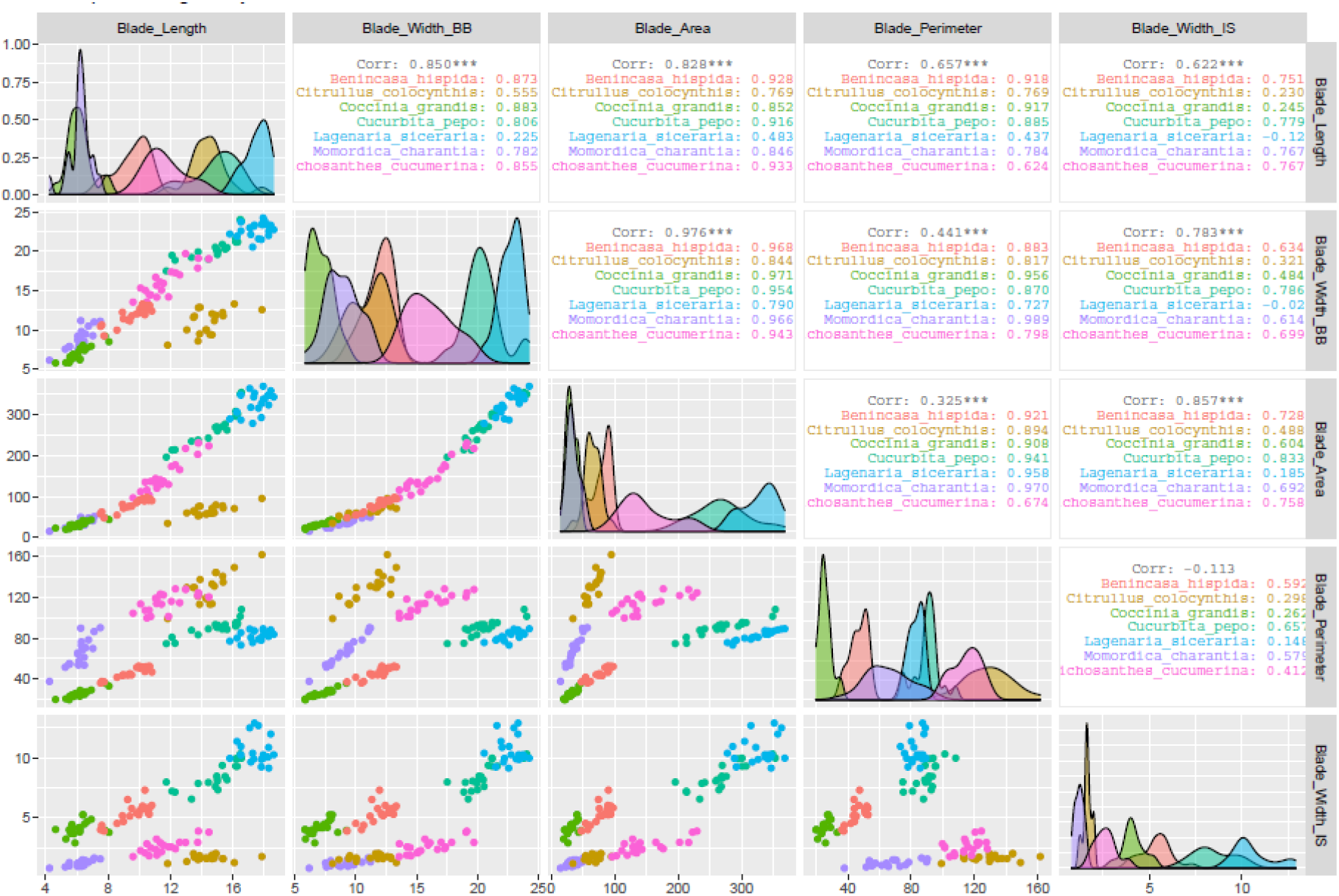
Scatterplot / Coefficient Matrix of the leaves’ blades dataset plotted with leaf blade variables only. The dataset excluding teeth variables show a positive correlation between all leaf blade variables within species and statistically significan differences between species.

There is an order of increasing length and width between all species with *C, grandis* and *M. charantia* both similar in length to *B. hispida*, *T. cucumerina*, *C. pepo*, *C. colocynthis*, and *L. siceraria*. The length measures the distance from the leaf apex to the leaf base. Just like the length, there are statistically significant differences between the width of all species, and variation is high.

The average leaf outlines of all the shapes studied is shown in (Fig. 1b). PCA for the leaf variables is show in Figure 4, and it indicates that most variations are observed in the first and second principal component are 76.2% and 19.8% respectively. PC1 and PC2 was further plotted against the leaf length (Fig. 2b, 2c), leaf width (Fig. 2d, 2e), and leaf perimeter (Fig. 2f, 2g) to determine the variations between the species. They were negatively correlated with the first and positively correlated with the second principal component.

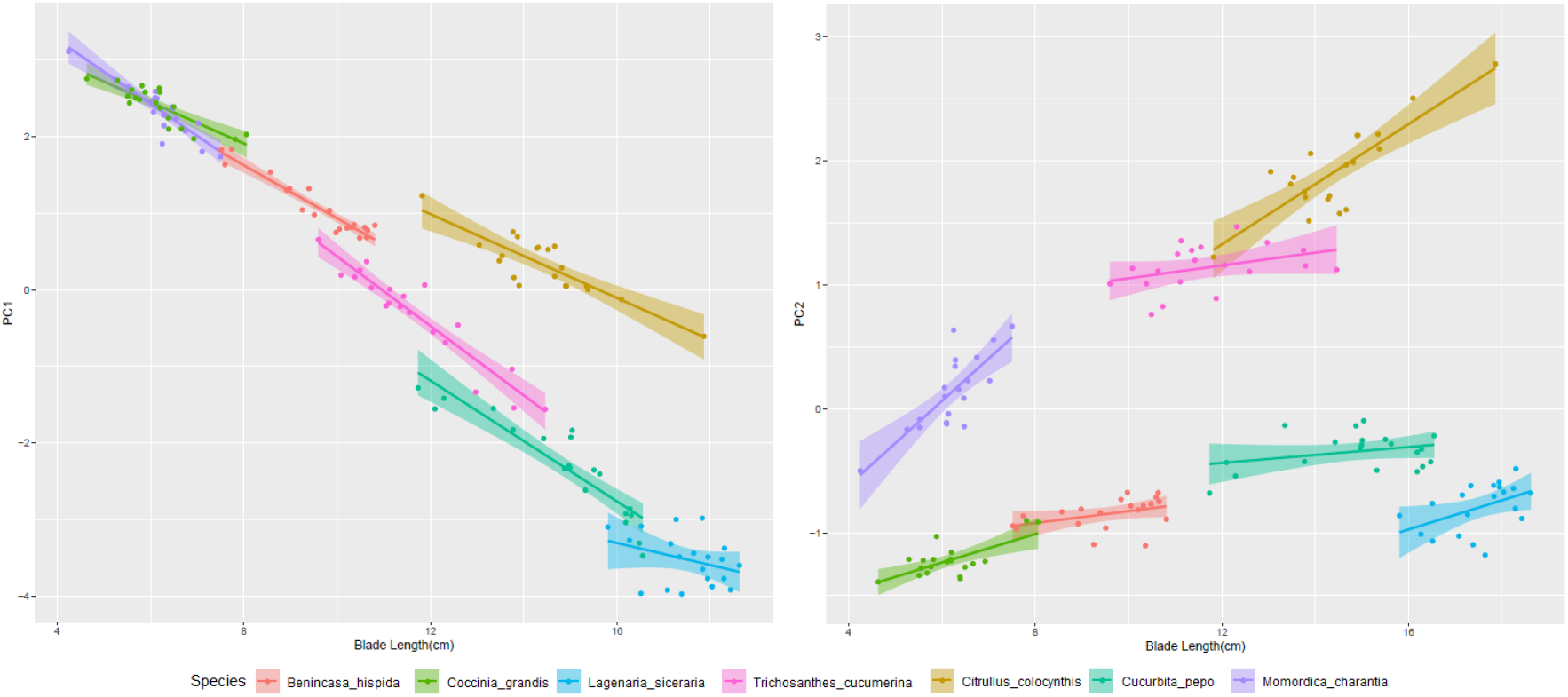
Fig. 2B (left), and Fig. 2C (right)

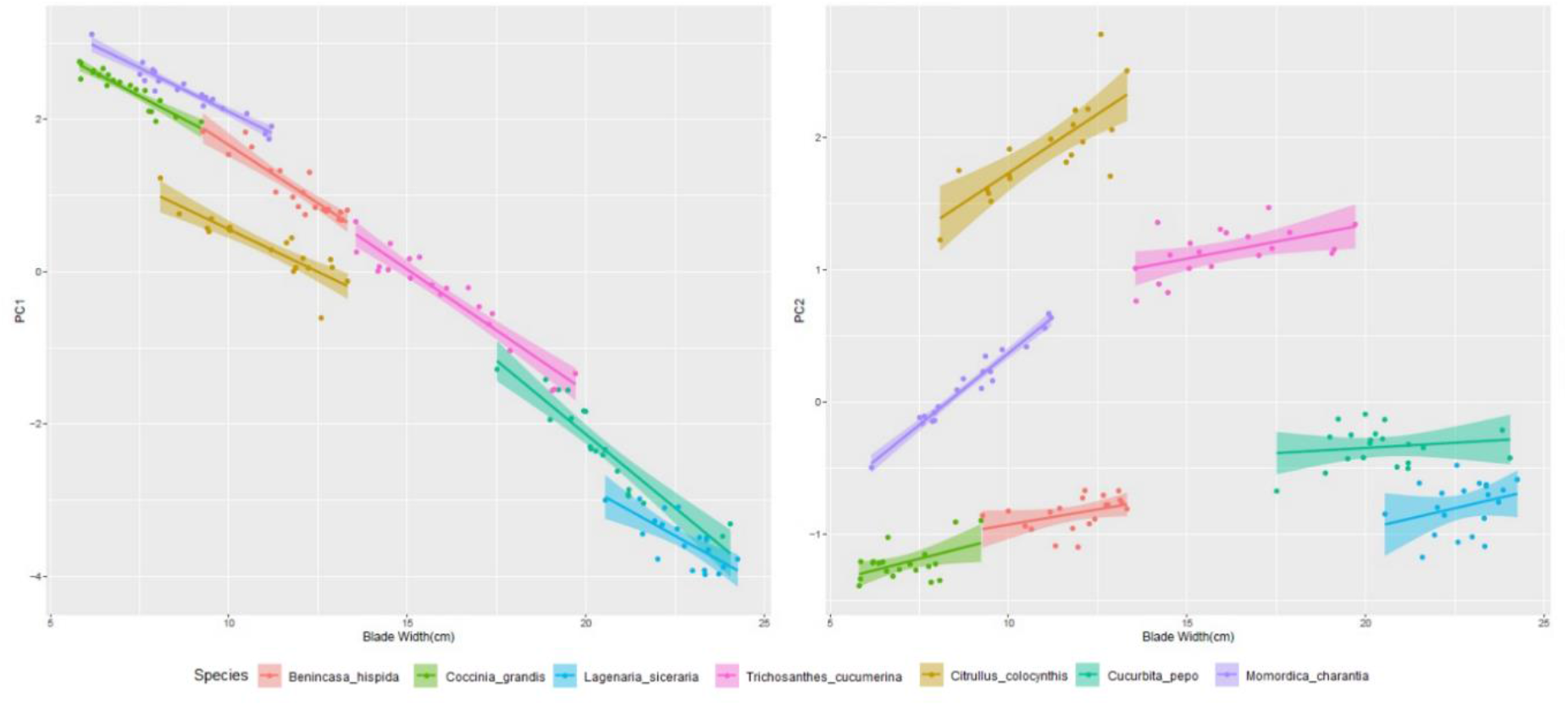
Fig. 2D (left) and 2E (right)

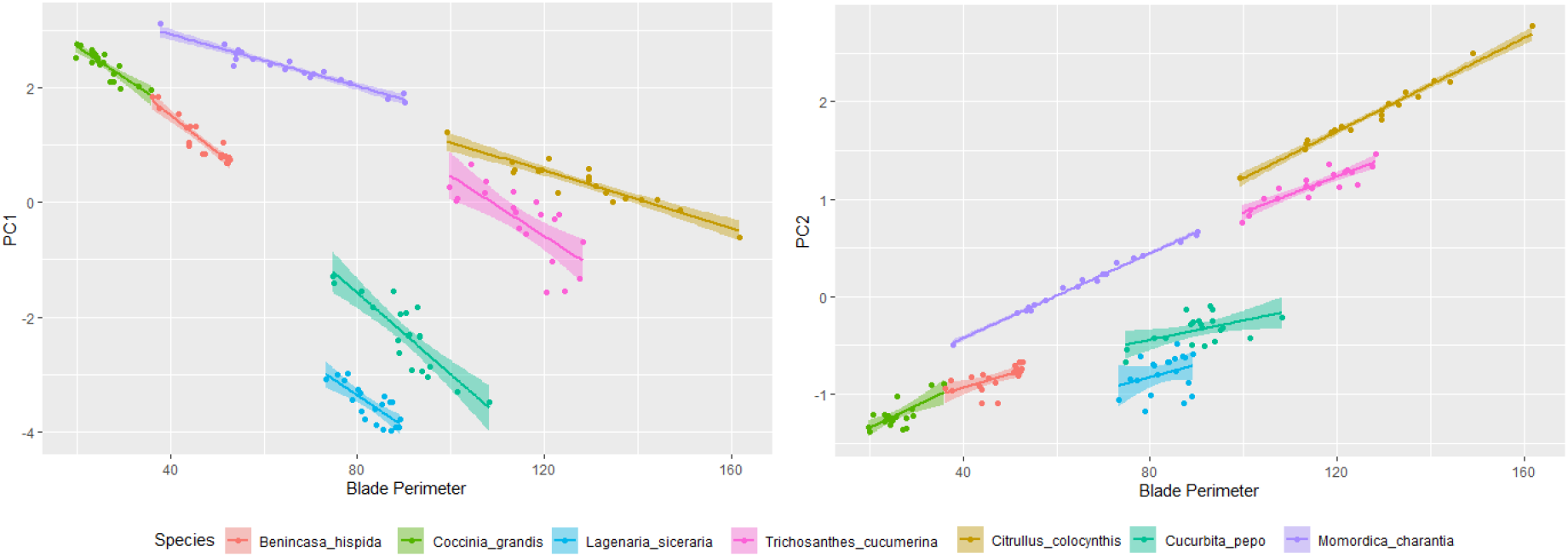
Fig. 2F (left) and 2G (right)

The area or total blade cover of the leaf another independently discriminative landmark of the leaf. Just as they share in the length and width, only *M. charantia* and *C. grandis* have an overlap in the leaf area (2 i).

It must be noted that this PCA are specifically for leaf variables, and all these variables altogether separate the dataset into seven morphospaces with no overlap. When considered individually, some of the variables overlapped in some species. *C. grandis* and *M. charantia* have the same length, *C. pepo* and *T. cucumerina* also have similar length.

The width-IS is an important informative landmark of the leaf. It represents the distance between the position 1 sinuses of both symmetry on the leaf (Biot et al., 2016). It showed a great degree of variation between each species with, and has no relationship between the length and width (2h).

Further discrimination between species, especially those that share overlap in landmarks, can be observed in perimeter (Fig. 2j). For example, the pair of *C. pepo* and *L. siceraria*, *T. cucumerina* and *C. colocynthis*, and *C. grandis* and *M. charantia*, each have similar area, but were separated by perimeter values (2k).

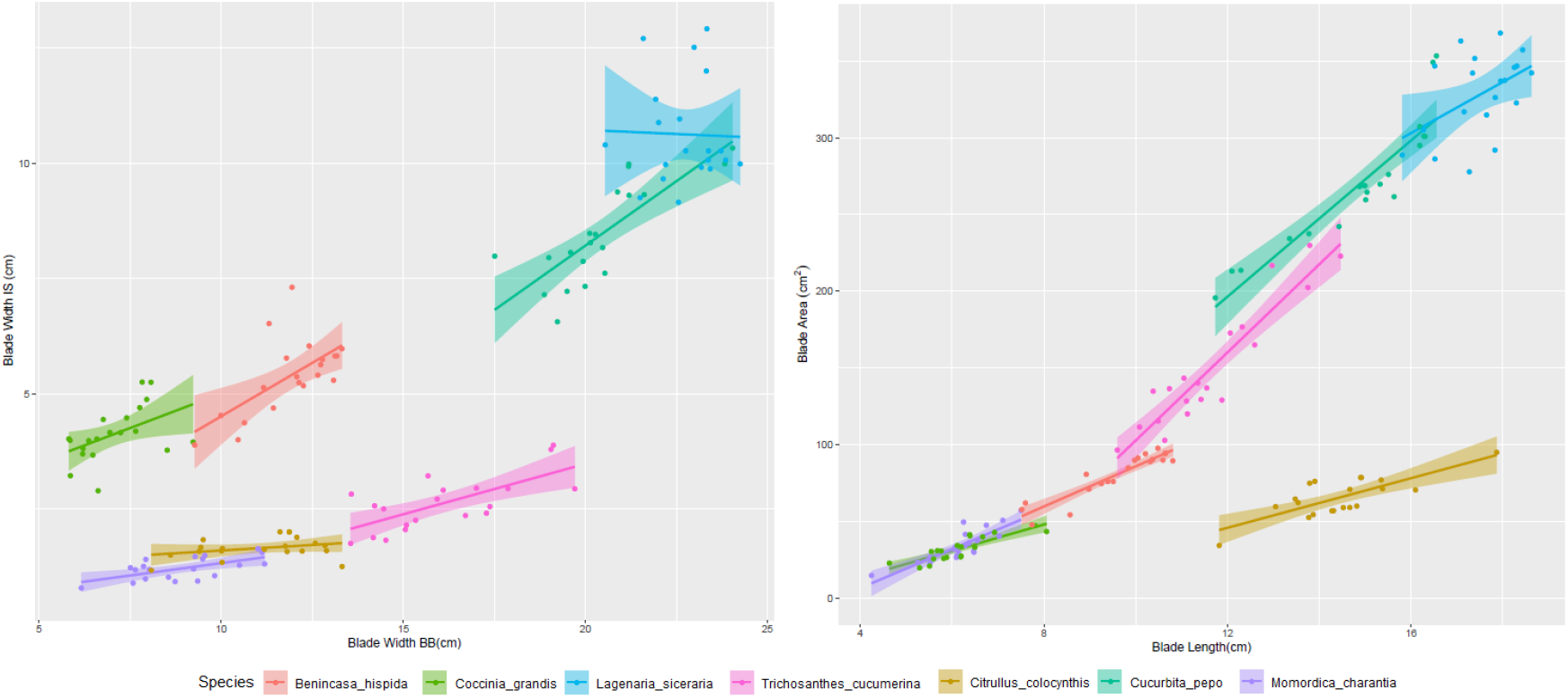
Fig. 2H (left) and 2I (right)

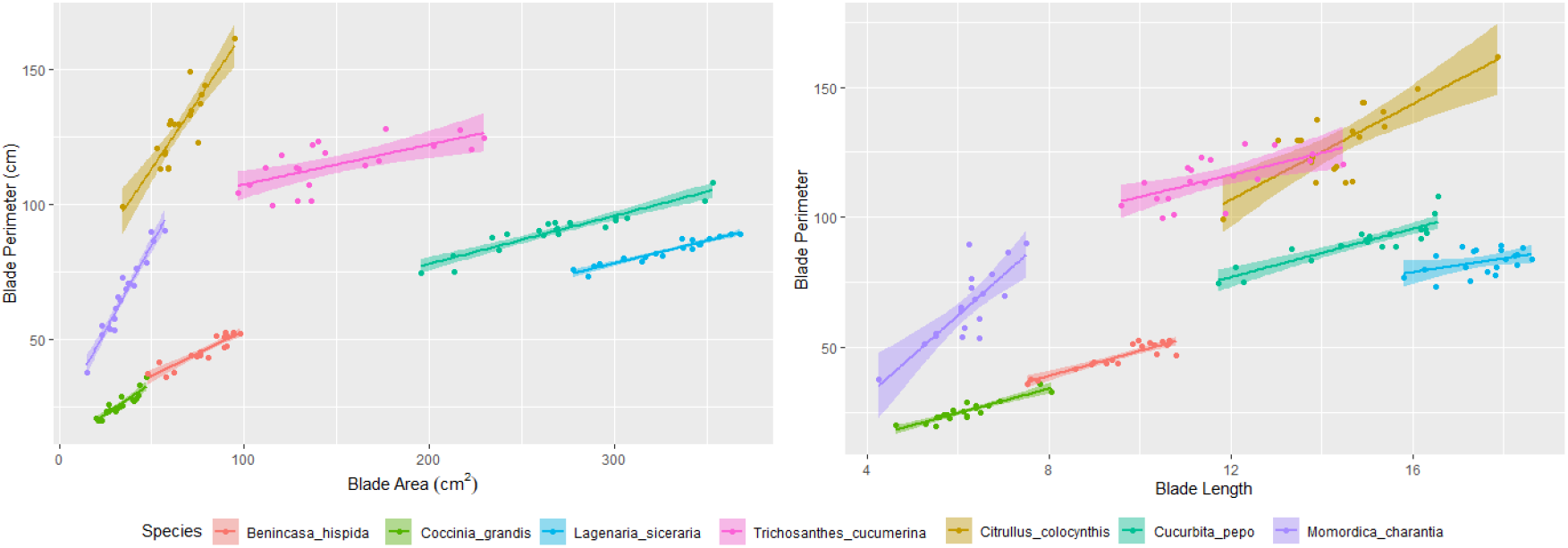
Fig. 2J and 2K

### Teeth Measures Analysis

Similar set of analysis carried out on the leaf dataset was carried out on the teeth landmarks dataset. The teeth dataset was further processed from 140 samples into 840 samples – total number of teeth per species multiplied by number of specimens per species, and this was done to treat each tooth position as an entity of its own (Supplementary 1 – *the leaf tip doesn’t count as teeth, it counts as apex*). The number of teeth varies between 4 and 10 (Fig. 3a), all showing unique teeth structure, and the total teeth number per species and the teeth position in relation to blade length varies across species. The teeth variables considered during PCA include teeth position, width, height altitude, height median, tooth area, perimeter, height of tooth position from base, and height of tooth position from tip.

**Fig. 3A:**
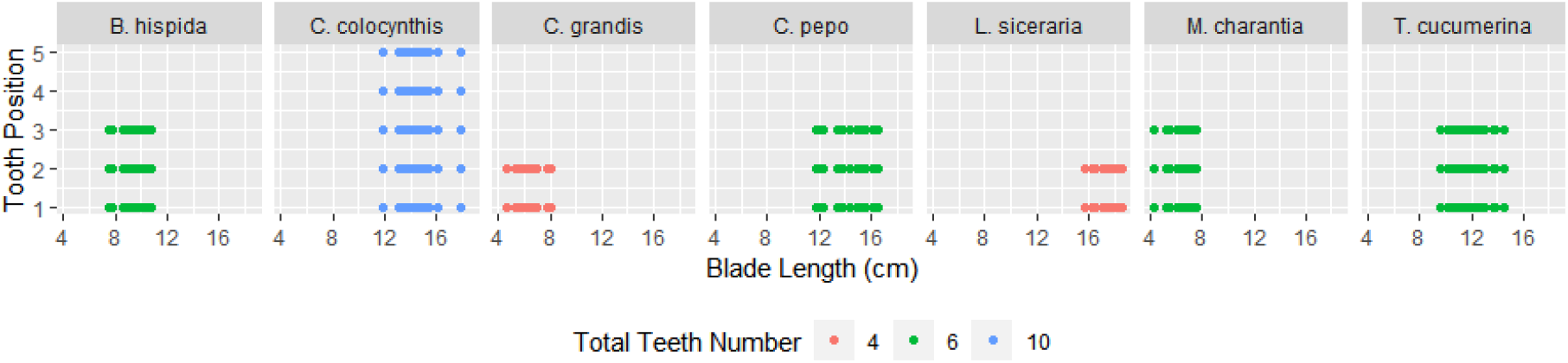
Total teeth number per species and the teeth position in relation to blade length varies across species.

Figure 3b and 3c shows a scatterplot of the first four PCs with all leaf and teeth variables, 13 in all, with PC1 responsible for the most variation. The inclusion of the teeth variable caused overlaps in the grouping which could be attributed the teeth positions on both sides of the blade symmetry. However, considering that the species were known apriori, they were still separated rightly and accordingly into seven groups, each one representing a species, with the tooth position then separating each species into different subgroup based on the number of species. The landmarks of the teeth were then subjected to a correlation coefficient matrix as shown in Fig. 3d and expanded in a pairwise scatterplot of all landmarks (supplementary 2).

**Fig. 3B:**
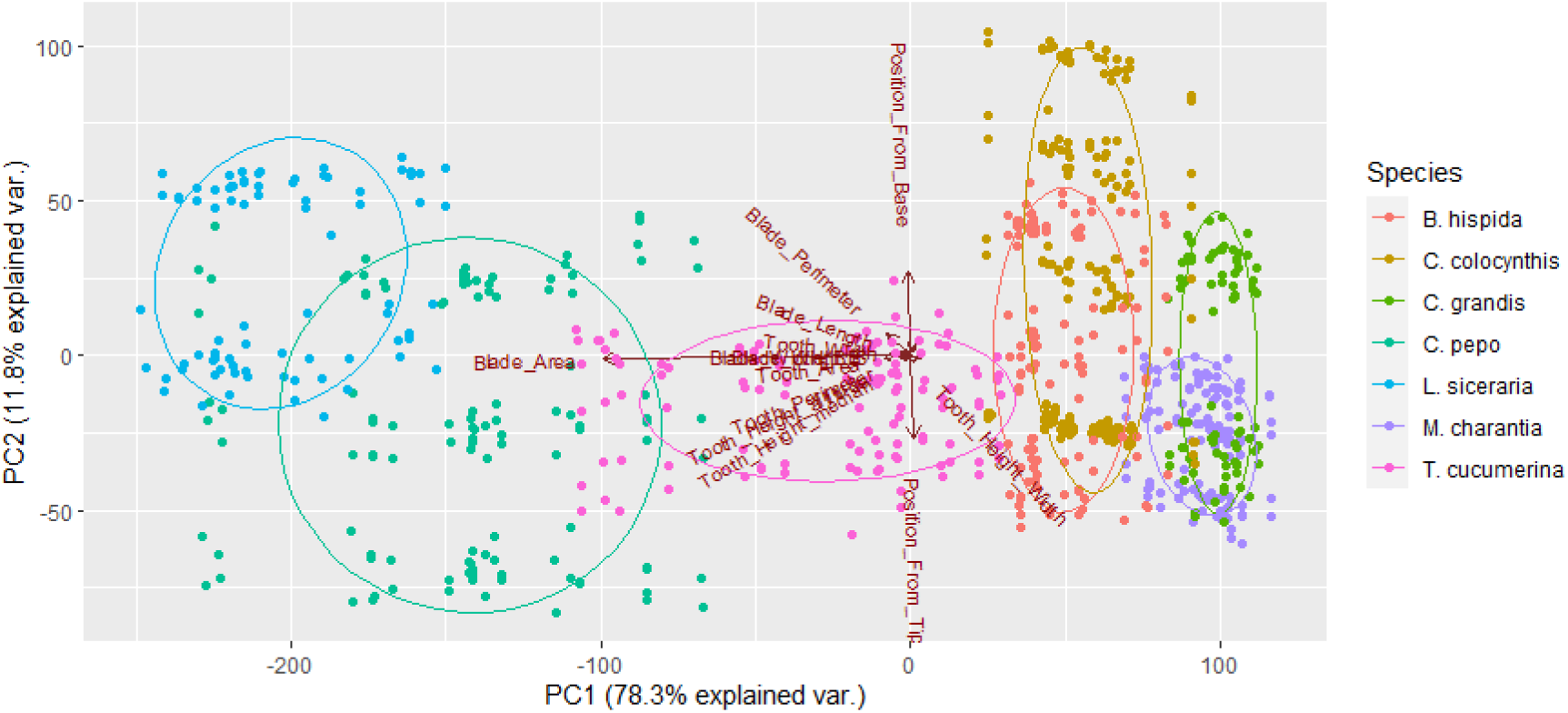
PCA of all dataset including teeth and blade. PC1 contributed the most to the variation within the group studied at 78.3%, while PC2 contributed 11.8%.

**Fig. 3C:**
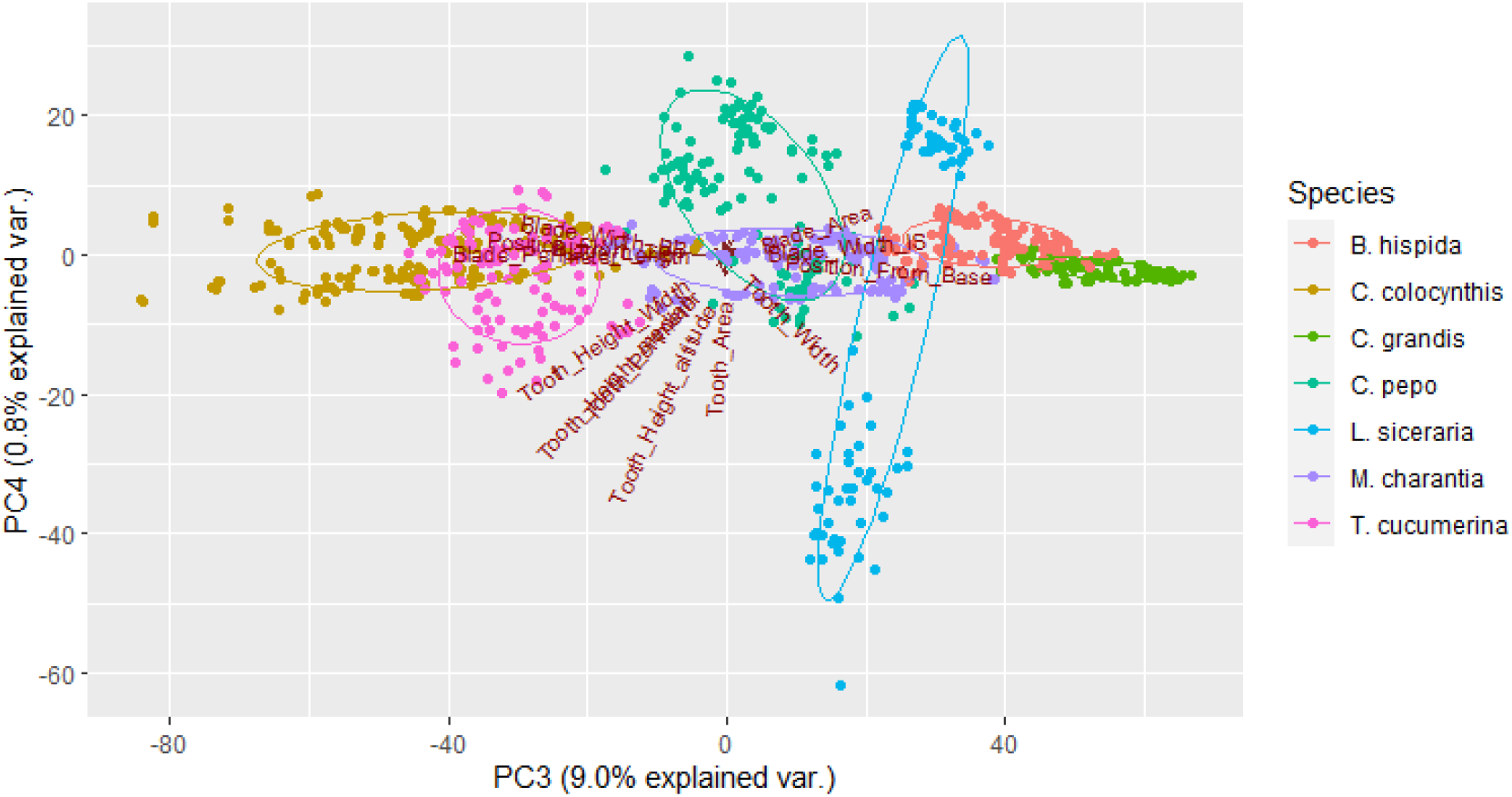
PCA of all dataset including teeth and blade. PC3 contributed the least of the three most important PCs at 9% to the variation.

**Fig. 3D:**
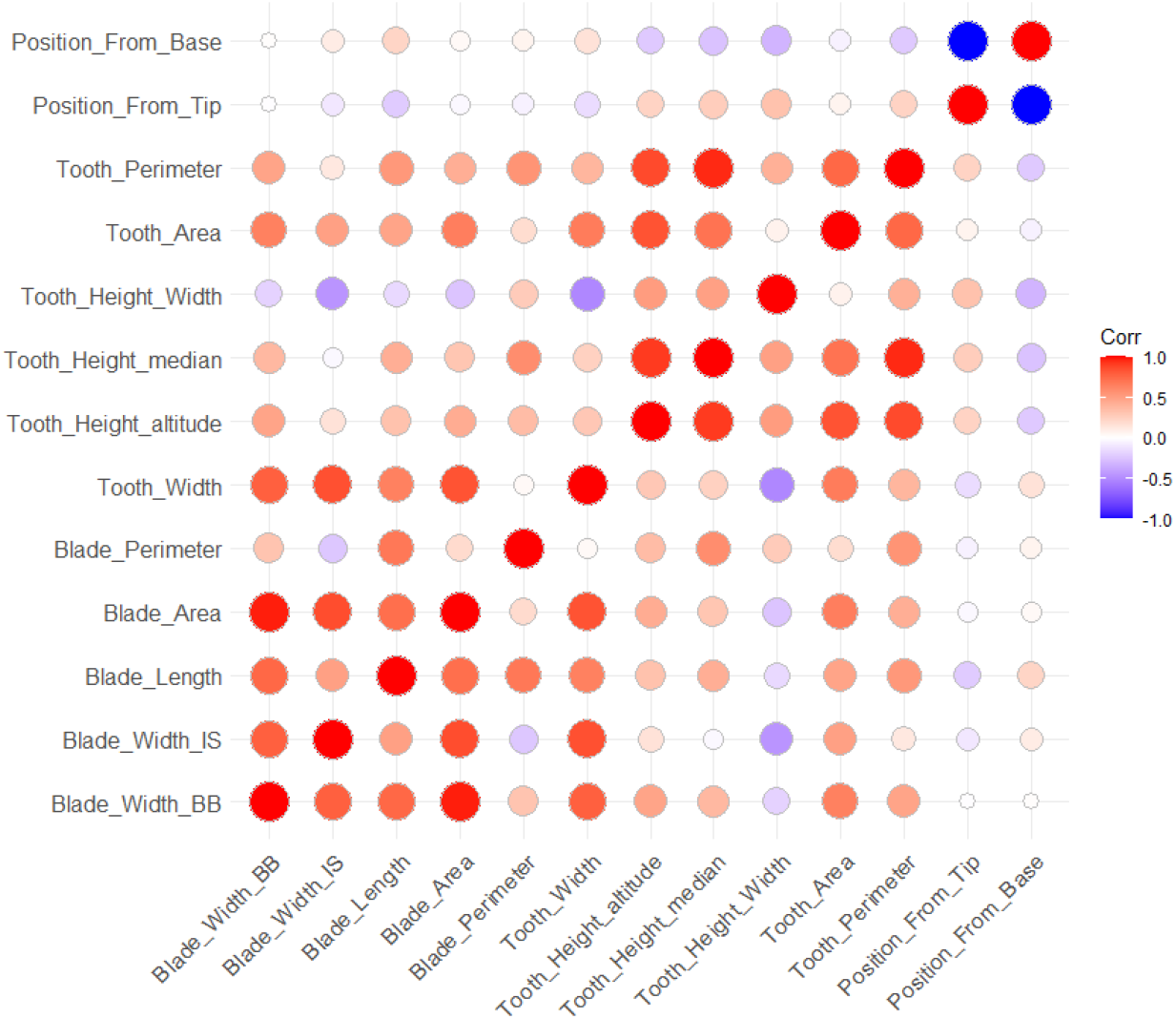
Correlation coefficient matrix of all teeth landmarks. From Negatively correlated variables (−1, blue color) to positively correlated variables (+1, red color).

The tooth position from tip, and the tooth position from base are the most negatively correlated because the move in opposite directions. The blade height to width ratio is also negatively correlated against the leaf variables except the leaf perimeter, because it contributes significantly to the proportion of leaf perimeter, but its contribution is insignificant to the height or width of the leaf.

The tooth perimeter of each tooth relative to the blade perimeter is a highly variable feature (Fig. 3e). While the teeth perimeter has low values on position one in some species, others have higher values on position one. The same applied to position 2, 3, 4, and 5, across all species. There’s also a positive correlation between the teeth perimeter and the blade perimeter – the higher the value of the blade perimeter, the higher the value of the teeth perimeter. Similar situation was observed in the tooth perimeter relative to the blade perimeter (Fig. 3f). This is a result of the differences in tooth size per position on the leaf. The tooth size (width & height) on both position of the symmetry isn’t too far apart in values (Fig. 3g), i.e. the tooth size on position 1 of the left symmetry is close to the size of the position 1 on the right symmetry, the same apply to position 2 on the left and position 2 on the right.

**Fig. 3E:**
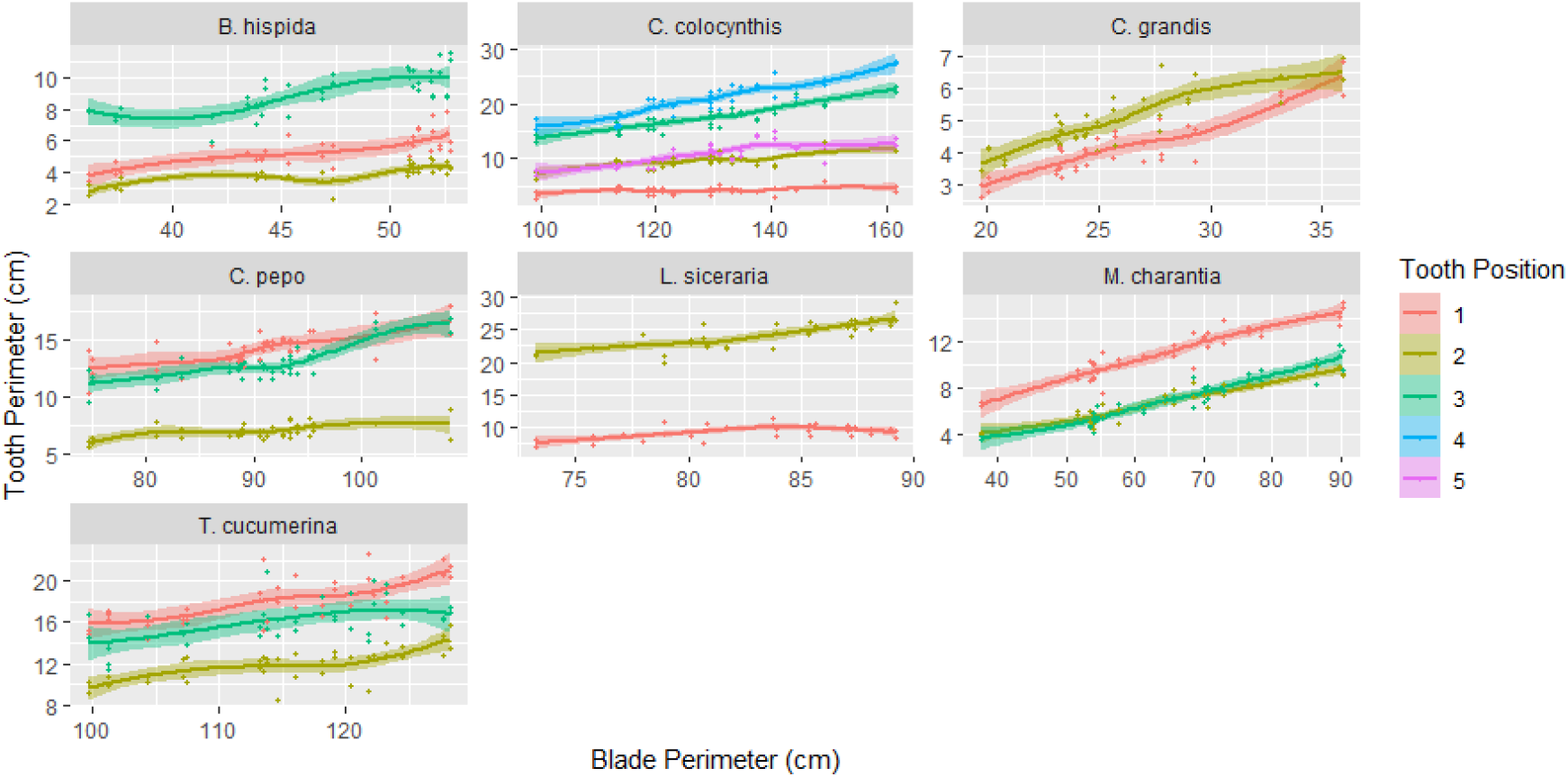
Tooth perimeter of each tooth relative to the blade perimeter is a highly variable landmark

**Fig. 3F:**
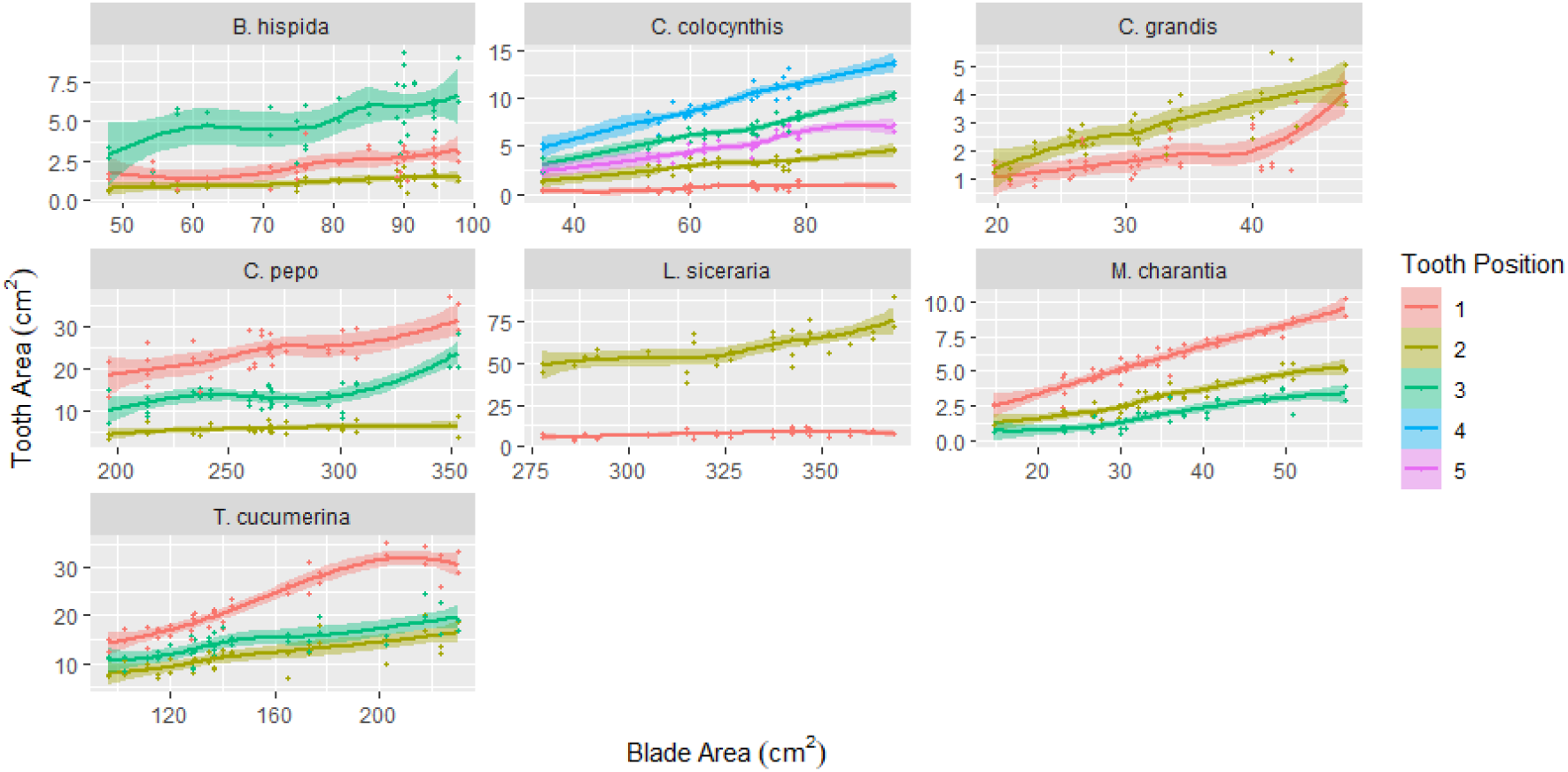
Tooth area contributes significantly to the blade area.

**Fig. 3G:**
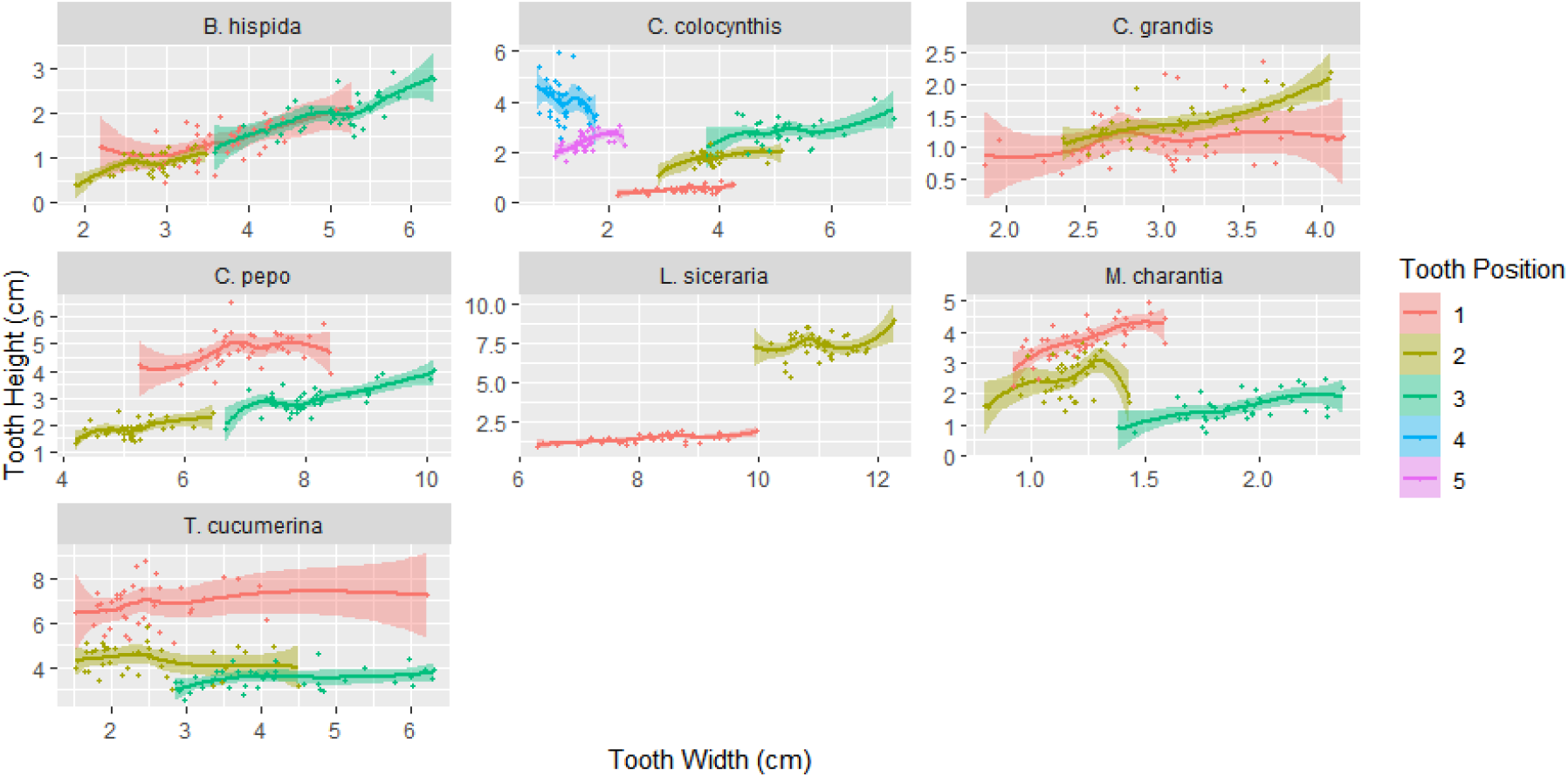
Tooth height and width varies per tooth position within and between each species

When all variables were plotted on a correlation matrix against PC1, PC2, and PC3, from an all-inclusive blade dataset (including the teeth landmarks), it shows important landmarks contributing to the variations in each of the PCs (Fig. 3h). The blade and tooth area are most important in contributing to variation in PC1, while blade perimeter and position of each of this tooth from tip and base were the most important landmarks contributing to variation in PC2. In PC3, the position of each of the tooth from tip and base, and blade and tooth perimeter were the landmarks that contributed the most to variations to PC3.

**Fig. 3H:**
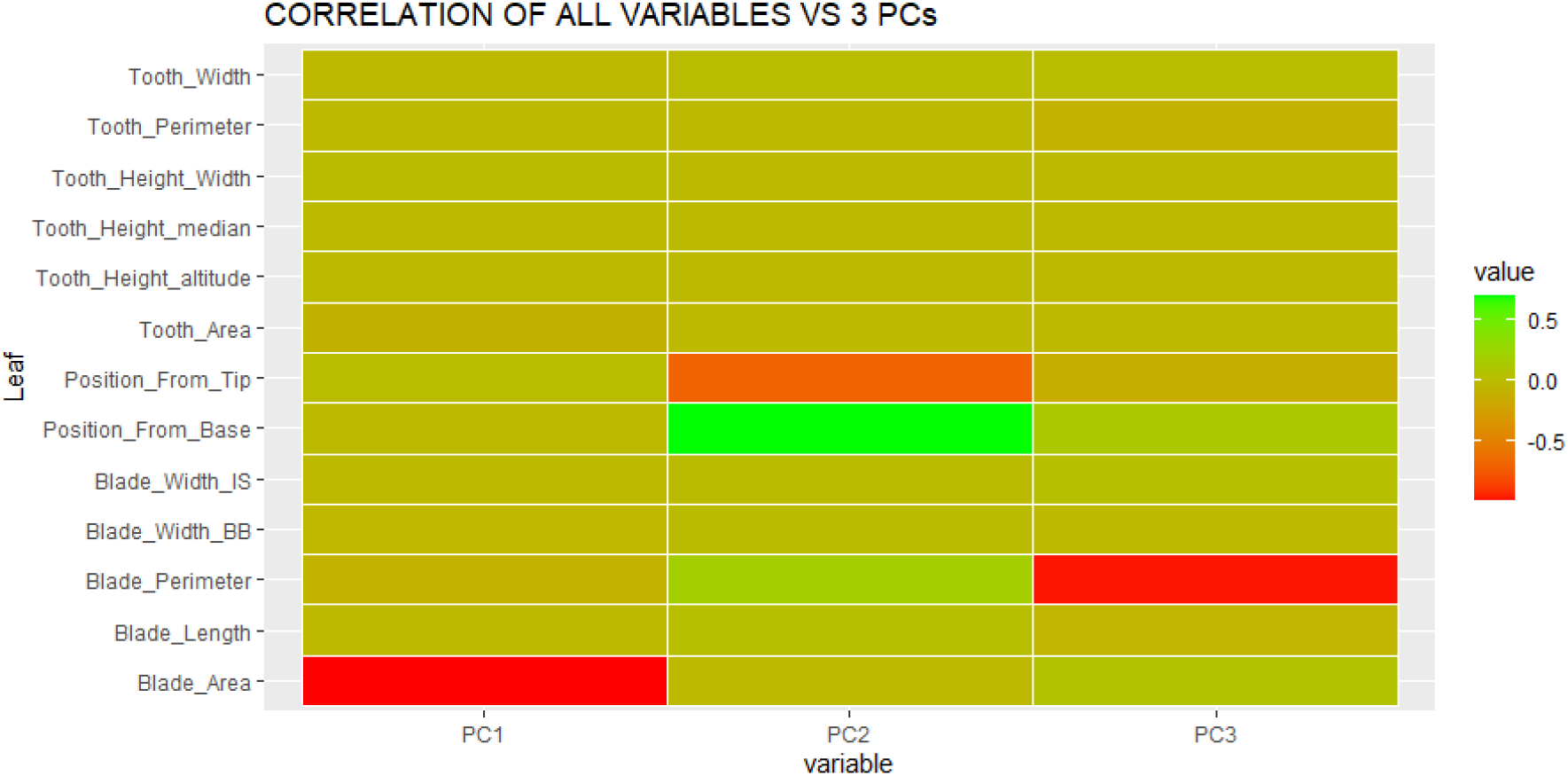

## DISCUSSION

There has been an evolution in the visualization and analysis of leaf shapes in moderns times from ImageJ (Abramoff et al., 2004) to LEAFPROCESSOR (Backhaus et al., 2010) MorphoJ (Klingenberg, 2011), MorphoLeaf (Biot et al., 2016), and MASS (Chuanromanee et al., 2019), with different scientists proving to have effectively applied them for almost similar objectives. These tools allow of ease of use and replicability in application. While others rely on a combination of multiple software for landmarking, outlining, and analysis, coupled with the import and export of file formats, Morpholeaf is a reliable software for landmark selection, in-software image analysis, superimposition, and visualization. It does it all as a plugin in FreeD (Andrey and Maurin, 2005), which comes with an S-viewer for 2D or 3D visualization of the shapes.

The seven species studied had leaf shapes ranging from simple to complex (Fig. 1a), and the shape variation was visibly separated into seven groups with all landmarks important in this grouping. When MorphoLeaf is applied to more species within the family, it would cluster the species into groups reliable enough to reveal the influence of leaf landmarks and teeth variations on evolution within the group. ImageJ (Abramoff et al., 2004), a similar tool similar tool for landmark selection has been used along with other applications for subsequent processing and analysis in identification and classification (Corney et al., 2012), genetic basis for Arabidopsis serration (Biot et al., 2016), and predicting species position and the role of leaf landmark in evolution and development in *Passiflora* (Chitwood and Otoni, 2017) and *Vitis* (Chitwood et al., 2016).

This work is an image-based inter-specific geomorphometric analysis (digital morphometry), and its potential in application its effectiveness to a larger group can be observed in the differentiation between the species. There different patterns in the leaf perimeters, leaf area, tooth area, tooth perimeter, shows that different node positions will be observed when applied to analysis within a larger group. The question remains - would a GMM-based clustering be grouped the same way as a gene-based phylogeny? Possibly yes, as some level of success was achieved in revealing the evolution of shapes within *Oxalis* when geomorphometric data was combined to molecular data (Morello et al., 2018). But would it contribute to a more robust hypothesis on evolutionary and functional significance? Definitely yes. The variations in leaf shape across plant groups either due to climatic variables or genetic modifications, with different expressions and sizes representing the effects of temperature, rainfall, exposure to sunlight, and mutations, have provided scientists reasons to hypothesize the evolutionary importance of leaf shape (Edwards et al., 2017; Gallaher et al., 2019; Kidner and Umbreen, 2010; Nicotra et al., 2011).

Previous morphometric studies in the family Cucurbitaceae have focuses on few species largely focusing on qualitative characters or measurements of characters, therefore outlining characters that are important but limited to identification of the species and morphological characterization of the varieties, but not for a robust delimitation of the super-taxa. Our analysis suggests that a PCA of the leaf outlines and teeth clearly segregates the species into seven groups as apriori determined. Each of the variables contributed to this separation, but the blade area, blade perimeter, tooth area, tooth perimeter, height of (each position of the) tooth from tip, and the height of each (position of the) tooth from base are important and informative factors when quantitatively determining the placement of each species into the correct group. This should however not be used in isolation, but in the same weight as traditionally used characters like leaf type, apex, base, venation, amongst other qualitative characters, in understanding evolution within a plant group. The relationship between vasculature and leaf blade is also of great importance (Chitwood and Otoni, 2017), as it contributes as much as 15 additional informative landmarks

Developmental studies on different genotypes (cultivars and varieties) of species have shown divergence from the basic shapes in heteroblastic situations which are usually a result of genetic or environmental factors (Chitwood et al., 2012; Ostria-Gallardo et al., 2016), therefore, intrinsic differences within each species cannot be ignored. And because the samples used in this study were collected from multiple populations, the different ecological variables from the environment they were collected from could be a responsible factor for these differences. This, however, doesn’t take away the base leaf outline of each species, it only further underscores the need for leaving out bias and taking all factors characters into consideration when tracking evolution within a plant group, or doing other bioinformatic works. The ease of use and efficiency of digital morphometrics should encourage botanists to draw more accurate inferences to from a comprehensive pool of data to support their answer their research questions (Luca and Annamaria, 2019).

There have already been comprehensive morphometric studies (Ekeke and Agogbua, 2018; Josephine et al., 2015), phylogenetic studies (Chomicki et al., 2019; Misra et al., 2017; Renner and Pandey, 2013; Schaefer and Renner, 2011; Zhang et al., 2006), and recently, phylogenomic studies (Guo et al., 2020) on Cucurbitaceae with none referring to the important characters listed earlier. The bias here is because Cucurbitaceae is used as a model in this study, the same however applies to most other researches in other literatures studied. Although digital morphometry only of recent beginning to reveal these additional characters, it is important that they are used in subsequent studies. Similarly, digital morphometrics reveals that applying general Procrustes analysis, linear discriminant analysis, and elliptical fourier descriptors to morphological data can improve the result of bioinformatic researches on inter-specific and intra-specific plant populations, particularly when genetic and environmental data are used (Klein et al., 2017; Migicovsky et al., 2018). GMM revealed differences in two varieties of *Escallona* (Morello and Sede, 2016)

Although MorphoLeaf was designed with intra-specific models, this study shows its success in inter-specific leaf shape analysis, this is further supported by evidence from GMM analyses of leaf variation in four species of *Quercus* that shows a strong correlation between leaf shape and taxonomy of *Quercus* (Viscosi et al., 2009b, 2009a). MorphoLeaf’s success in identifying intra-specific and inter-specific leaf shape variations, just like other applications earlier mentioned, allows evolutionary biologists to use complementary traits to identify and classify plant, even divergent plants that have been modified as a result of adaption-inducing variables like the environment. These tools are leading to a shift back into further exploration of image-based geomorphometry to provide phenotypic data that can supplement already existing morphometric and molecular data for phylogeny, and GMM analysis presents itself as vital in developing applications, guides, devices, and ultimately a central virtual herbarium for plant identification (Agarwal et al., 2006; Belhumeur et al., 2008; Cope et al., 2012; Jamil et al., 2015; Schlautman et al., 2020).

As the shift from traditional morphometrics to digital morphometrics driven by research questions continues to be generally adopted, more applications and technological solutions will be created with the intention of mass botanical digitization and solving problems related to identification of plants. This mass digitization provides natural history collectors, included but not limited to herbarium and field experts, with opportunities to mobilize data to explore different research hypotheses, solve scientific problems, reduce data gaps and biases, and for bioinformatic purposes (Beaman and Cellinese, 2012; Kattge et al., 2020; Lorieul et al., 2019; Soltis, 2017; Soltis et al., 2018). Subsequent works will focus on the application of MorphoLeaf and other applications on herbarium specimens and other leaf types, particularly complex leaves.

## SUPPLEMENTARY MATERIAL

The Data for this article is available online at: https://github.com/osooluwatobia/cucurbitaceae-morpholeaf.

## REFERENCES

Abramoff, M., Magalhaes, P., Ram, S., MD Abràmoff, P.M.S.R., 2004. Image processing with ImageJ. Biophotonics Int. 11, 36–43.

Adams, D.C., Otárola-Castillo, E., 2013. Geomorph: An r package for the collection and analysis of geometric morphometric shape data. Methods Ecol. Evol. 4, 393–399. https://doi.org/10.1111/2041-210X.12035

Agarwal, G., Belhumeur, P., Feiner, S., Jacobs, D., Kress, W.J., Ramamoorthi, R., Bourg, N.A., Dixit, N., Ling, H., Mahajan, D., Russell, R., Shirdhonkar, S., Sunkavalli, K., White, S., 2006. First steps toward an electronic field guide for plants. Taxon 55, 597–610. https://doi.org/10.2307/25065637

Alasadi, A.H.H., Anduljalil, E.Q., Khaleel, A.H., 2017. Leaf Recognition based on Neural Network Feed-Forward and Support Vector Machine Leaf Recognition based on Neural Network Feed-Forward and Support Vector Machine Classifiers 6, 92–99.

Andrey, P., Maurin, Y., 2005. Free-D: An integrated environment for three-dimensional reconstruction from serial sections. J. Neurosci. Methods 145, 233–244. https://doi.org/10.1016/j.jneumeth.2005.01.006

Backhaus, A., Kuwabara, A., Bauch, M., Monk, N., Sanguinetti, G., Fleming, A., 2010. Leafprocessor: A new leaf phenotyping tool using contour bending energy and shape cluster analysis. New Phytol. 187, 251–261. https://doi.org/10.1111/j.1469-8137.2010.03266.x

Beaman, R.S., Cellinese, N., 2012. Mass digitization of scientific collections: New opportunities to transform the use of biological specimens and underwrite biodiversity science. Zookeys. https://doi.org/10.3897/zookeys.209.3313

Belhumeur, P., Chen, D., Feiner, S., Jacobs, D., Kress, W., 2008. Searching the world’s herbaria: A system for visual identification of plant species. Comput. Vis. - ECCV 2008, Springer.

Biot, E., Cortizo, M., Burguet, J., Kiss, A., Oughou, M., Maugarny-Calès, A., Gonçalves, B., Adroher, B., Andrey, P., Boudaoud, A., Laufs, P., 2016. Multiscale quantification of morphodynamics: Morpholeaf software for 2D shape analysis. Dev. 143, 3417–3428. https://doi.org/10.1242/dev.134619

Borges, L.M., Reis, V.C., Izbicki, R., 2020. Schrödinger’s phenotypes: Herbarium specimens show two-dimensional images are both good and (not so) bad sources of morphological data. Methods Ecol. Evol. 11, 1296–1308. https://doi.org/10.1111/2041-210X.13450

Chitwood, D.H., Headland, L.R., Kumar, R., Peng, J., Maloof, J.N., Sinha, N.R., 2012. The developmental trajectory of leaflet morphology in wild tomato species. Plant Physiol. 158, 1230–1240. https://doi.org/10.1104/pp.111.192518

Chitwood, D.H., Klein, L.L., O’Hanlon, R., Chacko, S., Greg, M., Kitchen, C., Miller, A.J., Londo, J.P., 2016a. Latent developmental and evolutionary shapes embedded within the grapevine leaf. New Phytol. 210, 343–355. https://doi.org/10.1111/nph.13754

Chitwood, D.H., Otoni, W.C., 2017. Morphometric analysis of Passiflora leaves: The relationship between landmarks of the vasculature and elliptical Fourier descriptors of the blade. Gigascience 6, 1–13. https://doi.org/10.1093/gigascience/giw008

Chitwood, D.H., Rundell, S.M., Li, D.Y., Woodford, Q.L., Yu, T.T., Lopez, J.R., Greenblatt, D., Kang, J., Londo, J.P., 2016b. Climate and Developmental Plasticity: Interannual Variability in Grapevine Leaf Morphology. Plant Physiol. 170.

Chitwood, D.H., Sinha, N.R., 2016. Evolutionary and Environmental Forces Sculpting Leaf Development. Curr. Biol. 26, R297–R306. https://doi.org/10.1016/j.cub.2016.02.033

Chomicki, G., Schaefer, H., Renner, S.S., 2019. Origin and domestication of Cucurbitaceae crops: insights from phylogenies, genomics and archaeology. New Phytol. https://doi.org/10.1111/nph.16015

Chuanromanee, T.S., Cohen, J.I., Ryan, G.L., 2019. Morphological Analysis of Size and Shape (MASS): An integrative software program for morphometric analyses of leaves. Appl. Plant Sci. 7. https://doi.org/10.1002/aps3.11288

Cope, J.S.J., Corney, D.D., Clark, J.J.Y., Remagnino, P., Wilkin, P., 2012. Plant species identification using digital morphometrics: A review. Expert Syst. Appl. 39, 7562–7573. https://doi.org/10.1016/j.eswa.2012.01.073

Corney, D.P.A.A., Tang, H.L., Clark, J.Y., Hu, Y., Jin, J., 2012. Automating digital leaf measurement: The tooth, the whole tooth, and nothing but the tooth. PLoS One 7, 1–10. https://doi.org/10.1371/journal.pone.0042112

Dkhar, J., Pareek, A., 2014. What determines a leaf’s shape? Evodevo 5. https://doi.org/10.1186/2041-9139-5-47

Edwards, E.J., Chatelet, D.S., Spriggs, E.L., Johnson, E.S., Schlutius, C., Donoghue, M.J., 2017. Correlation, causation, and the evolution of leaf teeth: A reply to Givnish and Kriebel. Am. J. Bot. 104, 509–515. https://doi.org/10.3732/ajb.1700075

Edwards, E.J., Spriggs, E.L., Chatelet, D.S., Donoghue, M.J., 2016. Unpacking a century-old mystery: Winter buds and the latitudinal gradient in leaf form. Am. J. Bot. 103, 975–978. https://doi.org/10.3732/ajb.1600129

Ekeke, C., Agogbua, J.U., 2018. Morphological and Anatomical Studies on Trichosanthes cucumerina L. (Cucurbitaceae). Int. J. Plant Soil Sci. 25, 1–8. https://doi.org/10.9734/ijpss/2018/44982

Fu, G., Dai, X., Symanzik, J., Bushman, S., 2017. Quantitative gene–gene and gene– environment mapping for leaf shape variation using tree-based models. New Phytol. 213, 455–469. https://doi.org/10.1111/nph.14131

Gallaher, T.J., Adams, D.C., Attigala, L., Burke, S. V., Craine, J.M., Duvall, M.R., Klahs, P.C., Sherratt, E., Wysocki, W.P., Clark, L.G., 2019. Leaf shape and size track habitat transitions across forest–grassland boundaries in the grass family (Poaceae). Evolution (N. Y). 73, 927–946. https://doi.org/10.1111/evo.13722

Guo, J., Xu, W., Hu, Y., Huang, J., Zhao, Y., Zhang, L., Huang, C.H., Ma, H., 2020. Phylotranscriptomics in Cucurbitaceae Reveal Multiple Whole-Genome Duplications and Key Morphological and Molecular Innovations. Mol. Plant. https://doi.org/10.1016/j.molp.2020.05.011

Hernández-Esquivel, K.B., Piedra-Malagón, E.M., Cornejo-Tenorio, G., Mendoza-Cuenca, L., González-Rodríguez, A., Ruiz-Sanchez, E., Ibarra-Manríquez, G., 2020. Unraveling the extreme morphological variation in the neotropical Ficus aurea complex (subg. Spherosuke, sect. Americanae, Moraceae). J. Syst. Evol. 58, 263–281. https://doi.org/10.1111/jse.12564

Ichihashi, Y., Aguilar-Martínez, J.A., Farhi, M., Chitwood, D.H., Kumar, R., Millon, L. V., Peng, J., Maloof, J.N., Sinha, N.R., 2014. Evolutionary developmental transcriptomics reveals a gene network module regulating interspecific diversity in plant leaf shape. Proc. Natl. Acad. Sci. U. S. A. 111. https://doi.org/10.1073/pnas.1402835111

Itgen, M.W., Sessions, S.K., Wilson, L.D., Townsend, J.H., 2019. Integrative Systematic Revision of Bolitoglossa celaque (Caudata: Plethodontidae), with a new species from the Lenca Highlands of Honduras. Herpetol. Monogr. 33, 48–70. https://doi.org/10.1655/HERPMONOGRAPHS-D-19-00001.1

Jamil, N., Hussin, N.A.C., Nordin, S., Awang, K., 2015. Automatic Plant Identification: Is Shape the Key Feature? Procedia Comput. Sci. 76, 436–442. https://doi.org/10.1016/j.procs.2015.12.287

Josephine, A., Chimezie, E., Bosa, E.O., 2015. Morpho-anatomical characters of Zehneria capillacea (Schumach) C. Jeffrey and Zehneria scabra (L.F.) Sond Cucurbitaceae. African J. Plant Sci. 9, 457–465. https://doi.org/10.5897/ajps2015.1306

Kattge, J., Bönisch, G., Díaz, S., Lavorel, S., Prentice, I.C., Leadley, P., Tautenhahn, S., Werner, G.D.A., Aakala, T., Abedi, M., Acosta, A.T.R., Adamidis, G.C., Adamson, K., Aiba, M., Albert, C.H., Alcántara, J.M., Alcázar C. C., Aleixo, I., Ali, H., Amiaud, B., Ammer, C., Amoroso, M.M., Anand, M., Anderson, C., Anten, N., Antos, J., Apgaua, D.M.G., Ashman, T.L., Asmara, D.H., Asner, G.P., Aspinwall, M., Atkin, O., Aubin, I., Baastrup-Spohr, L., Bahalkeh, K., Bahn, M., Baker, T., Baker, W.J., Bakker, J.P., Baldocchi, D., Baltzer, J., Banerjee, A., Baranger, A., Barlow, J., Barneche, D.R., Baruch, Z., Bastianelli, D., Battles, J., Bauerle, W., Bauters, M., Bazzato, E., Beckmann, M., Beeckman, H., Beierkuhnlein, C., Bekker, R., Belfry, G., Belluau, M., Beloiu, M., Benavides, R., Benomar, L., Berdugo-Lattke, M.L., Berenguer, E., Bergamin, R., Bergmann, J., Bergmann Carlucci, M., Berner, L., Bernhardt-Römermann, M., Bigler, C., Bjorkman, A.D., Blackman, C., Blanco, C., Blonder, B., Blumenthal, D., Bocanegra-González, K.T., Boeckx, P., Bohlman, S., Böhning-Gaese, K., Boisvert-Marsh, L., Bond, W., Bond-Lamberty, B., Boom, A., Boonman, C.C.F., Bordin, K., Boughton, E.H., Boukili, V., Bowman, D.M.J.S., Bravo, S., Brendel, M.R., Broadley, M.R., Brown, K.A., Bruelheide, H., Brumnich, F., Bruun, H.H., Bruy, D., Buchanan, S.W., Bucher, S.F., Buchmann, N., Buitenwerf, R., Bunker, D.E., Bürger, J., Burrascano, S., Burslem, D.F.R.P., Butterfield, B.J., Byun, C., Marques, M., Scalon, M.C., Caccianiga, M., Cadotte, M., Cailleret, M., Camac, J., Camarero, J.J., Campany, C., Campetella, G., Campos, J.A., Cano-Arboleda, L., Canullo, R., Carbognani, M., Carvalho, F., Casanoves, F., Castagneyrol, B., Catford, J.A., Cavender-Bares, J., Cerabolini, B.E.L., Cervellini, M., Chacón-Madrigal, E., Chapin, K., Chapin, F.S., Chelli, S., Chen, S.C., Chen, A., Cherubini, P., Chianucci, F., Choat, B., Chung, K.S., Chytrý, M., Ciccarelli, D., Coll, L., Collins, C.G., Conti, L., Coomes, D., Cornelissen, J.H.C., Cornwell, W.K., Corona, P., Coyea, M., Craine, J., Craven, D., Cromsigt, J.P.G.M., Csecserits, A., Cufar, K., Cuntz, M., da Silva, A.C., Dahlin, K.M., Dainese, M., Dalke, I., Dalle Fratte, M., Dang-Le, A.T., Danihelka, J., Dannoura, M., Dawson, S., de Beer, A.J., De Frutos, A., De Long, J.R., Dechant, B., Delagrange, S., Delpierre, N., Derroire, G., Dias, A.S., Diaz-Toribio, M.H., Dimitrakopoulos, P.G., Dobrowolski, M., Doktor, D., Dřevojan, P., Dong, N., Dransfield, J., Dressler, S., Duarte, L., Ducouret, E., Dullinger, S., Durka, W., Duursma, R., Dymova, O., E-Vojtkó, A., Eckstein, R.L., Ejtehadi, H., Elser, J., Emilio, T., Engemann, K., Erfanian, M.B., Erfmeier, A., Esquivel-Muelbert, A., Esser, G., Estiarte, M., Domingues, T.F., Fagan, W.F., Fagúndez, J., Falster, D.S., Fan, Y., Fang, J., Farris, E., Fazlioglu, F., Feng, Y., Fernandez-Mendez, F., Ferrara, C., Ferreira, J., Fidelis, A., Finegan, B., Firn, J., Flowers, T.J., Flynn, D.F.B., Fontana, V., Forey, E., Forgiarini, C., François, L., Frangipani, M., Frank, D., Frenette-Dussault, C., Freschet, G.T., Fry, E.L., Fyllas, N.M., Mazzochini, G.G., Gachet, S., Gallagher, R., Ganade, G., Ganga, F., García-Palacios, P., Gargaglione, V., Garnier, E., Garrido, J.L., de Gasper, A.L., Gea-Izquierdo, G., Gibson, D., Gillison, A.N., Giroldo, A., Glasenhardt, M.C., Gleason, S., Gliesch, M., Goldberg, E., Göldel, B., Gonzalez-Akre, E., Gonzalez-Andujar, J.L., González-Melo, A., González-Robles, A., Graae, B.J., Granda, E., Graves, S., Green, W.A., Gregor, T., Gross, N., Guerin, G.R., Günther, A., Gutiérrez, A.G., Haddock, L., Haines, A., Hall, J., Hambuckers, A., Han, W., Harrison, S.P., Hattingh, W., Hawes, J.E., He, T., He, P., Heberling, J.M., Helm, A., Hempel, S., Hentschel, J., Hérault, B., Hereş, A.M., Herz, K., Heuertz, M., Hickler, T., Hietz, P., Higuchi, P., Hipp, A.L., Hirons, A., Hock, M., Hogan, J.A., Holl, K., Honnay, O., Hornstein, D., Hou, E., Hough-Snee, N., Hovstad, K.A., Ichie, T., Igić, B., Illa, E., Isaac, M., Ishihara, M., Ivanov, L., Ivanova, L., Iversen, C.M., Izquierdo, J., Jackson, R.B., Jackson, B., Jactel, H., Jagodzinski, A.M., Jandt, U., Jansen, S., Jenkins, T., Jentsch, A., Jespersen, J.R.P., Jiang, G.F., Johansen, J.L., Johnson, D., Jokela, E.J., Joly, C.A., Jordan, G.J., Joseph, G.S., Junaedi, D., Junker, R.R., Justes, E., Kabzems, R., Kane, J., Kaplan, Z., Kattenborn, T., Kavelenova, L., Kearsley, E., Kempel, A., Kenzo, T., Kerkhoff, A., Khalil, M.I., Kinlock, N.L., Kissling, W.D., Kitajima, K., Kitzberger, T., Kjøller, R., Klein, T., Kleyer, M., Klimešová, J., Klipel, J., Kloeppel, B., Klotz, S., Knops, J.M.H., Kohyama, T., Koike, F., Kollmann, J., Komac, B., Komatsu, K., König, C., Kraft, N.J.B., Kramer, K., Kreft, H., Kühn, I., Kumarathunge, D., Kuppler, J., Kurokawa, H., Kurosawa, Y., Kuyah, S., Laclau, J.P., Lafleur, B., Lallai, E., Lamb, E., Lamprecht, A., Larkin, D.J., Laughlin, D., Le Bagousse-Pinguet, Y., le Maire, G., le Roux, P.C., le Roux, E., Lee, T., Lens, F., Lewis, S.L., Lhotsky, B., Li, Y., Li, X., Lichstein, J.W., Liebergesell, M., Lim, J.Y., Lin, Y.S., Linares, J.C., Liu, C., Liu, D., Liu, U., Livingstone, S., Llusià, J., Lohbeck, M., López-García, Á., Lopez-Gonzalez, G., Lososová, Z., Louault, F., Lukács, B.A., Lukeš, P., Luo, Y., Lussu, M., Ma, S., Maciel Rabelo Pereira, C., Mack, M., Maire, V., Mäkelä, A., Mäkinen, H., Malhado, A.C.M., Mallik, A., Manning, P., Manzoni, S., Marchetti, Z., Marchino, L., Marcilio-Silva, V., Marcon, E., Marignani, M., Markesteijn, L., Martin, A., Martínez-Garza, C., Martínez-Vilalta, J., Mašková, T., Mason, K., Mason, N., Massad, T.J., Masse, J., Mayrose, I., McCarthy, J., McCormack, M.L., McCulloh, K., McFadden, I.R., McGill, B.J., McPartland, M.Y., Medeiros, J.S., Medlyn, B., Meerts, P., Mehrabi, Z., Meir, P., Melo, F.P.L., Mencuccini, M., Meredieu, C., Messier, J., Mészáros, I., Metsaranta, J., Michaletz, S.T., Michelaki, C., Migalina, S., Milla, R., Miller, J.E.D., Minden, V., Ming, R., Mokany, K., Moles, A.T., Molnár, A., Molofsky, J., Molz, M., Montgomery, R.A., Monty, A., Moravcová, L., Moreno-Martínez, A., Moretti, M., Mori, A.S., Mori, S., Morris, D., Morrison, J., Mucina, L., Mueller, S., Muir, C.D., Müller, S.C., Munoz, F., Myers-Smith, I.H., Myster, R.W., Nagano, M., Naidu, S., Narayanan, A., Natesan, B., Negoita, L., Nelson, A.S., Neuschulz, E.L., Ni, J., Niedrist, G., Nieto, J., Niinemets, Ü., Nolan, R., Nottebrock, H., Nouvellon, Y., Novakovskiy, A., Nystuen, K.O., O’Grady, A., O’Hara, K., O’Reilly-Nugent, A., Oakley, S., Oberhuber, W., Ohtsuka, T., Oliveira, R., Öllerer, K., Olson, M.E., Onipchenko, V., Onoda, Y., Onstein, R.E., Ordonez, J.C., Osada, N., Ostonen, I., Ottaviani, G., Otto, S., Overbeck, G.E., Ozinga, W.A., Pahl, A.T., Paine, C.E.T., Pakeman, R.J., Papageorgiou, A.C., Parfionova, E., Pärtel, M., Patacca, M., Paula, S., Paule, J., Pauli, H., Pausas, J.G., Peco, B., Penuelas, J., Perea, A., Peri, P.L., Petisco-Souza, A.C., Petraglia, A., Petritan, A.M., Phillips, O.L., Pierce, S., Pillar, V.D., Pisek, J., Pomogaybin, A., Poorter, H., Portsmuth, A., Poschlod, P., Potvin, C., Pounds, D., Powell, A.S., Power, S.A., Prinzing, A., Puglielli, G., Pyšek, P., Raevel, V., Rammig, A., Ransijn, J., Ray, C.A., Reich, P.B., Reichstein, M., Reid, D.E.B., Réjou-Méchain, M., de Dios, V.R., Ribeiro, S., Richardson, S., Riibak, K., Rillig, M.C., Riviera, F., Robert, E.M.R., Roberts, S., Robroek, B., Roddy, A., Rodrigues, A.V., Rogers, A., Rollinson, E., Rolo, V., Römermann, C., Ronzhina, D., Roscher, C., Rosell, J.A., Rosenfield, M.F., Rossi, C., Roy, D.B., Royer-Tardif, S., Rüger, N., Ruiz-Peinado, R., Rumpf, S.B., Rusch, G.M., Ryo, M., Sack, L., Saldaña, A., Salgado-Negret, B., Salguero-Gomez, R., Santa-Regina, I., Santacruz-García, A.C., Santos, J., Sardans, J., Schamp, B., Scherer-Lorenzen, M., Schleuning, M., Schmid, B., Schmidt, M., Schmitt, S., Schneider, J. V., Schowanek, S.D., Schrader, J., Schrodt, F., Schuldt, B., Schurr, F., Selaya Garvizu, G., Semchenko, M., Seymour, C., Sfair, J.C., Sharpe, J.M., Sheppard, C.S., Sheremetiev, S., Shiodera, S., Shipley, B., Shovon, T.A., Siebenkäs, A., Sierra, C., Silva, V., Silva, M., Sitzia, T., Sjöman, H., Slot, M., Smith, N.G., Sodhi, D., Soltis, P., Soltis, D., Somers, B., Sonnier, G., Sørensen, M.V., Sosinski, E.E., Soudzilovskaia, N.A., Souza, A.F., Spasojevic, M., Sperandii, M.G., Stan, A.B., Stegen, J., Steinbauer, K., Stephan, J.G., Sterck, F., Stojanovic, D.B., Strydom, T., Suarez, M.L., Svenning, J.C., Svitková, I., Svitok, M., Svoboda, M., Swaine, E., Swenson, N., Tabarelli, M., Takagi, K., Tappeiner, U., Tarifa, R., Tauugourdeau, S., Tavsanoglu, C., te Beest, M., Tedersoo, L., Thiffault, N., Thom, D., Thomas, E., Thompson, K., Thornton, P.E., Thuiller, W., Tichý, L., Tissue, D., Tjoelker, M.G., Tng, D.Y.P., Tobias, J., Török, P., Tarin, T., Torres-Ruiz, J.M., Tóthmérész, B., Treurnicht, M., Trivellone, V., Trolliet, F., Trotsiuk, V., Tsakalos, J.L., Tsiripidis, I., Tysklind, N., Umehara, T., Usoltsev, V., Vadeboncoeur, M., Vaezi, J., Valladares, F., Vamosi, J., van Bodegom, P.M., van Breugel, M., Van Cleemput, E., van de Weg, M., van der Merwe, S., van der Plas, F., van der Sande, M.T., van Kleunen, M., Van Meerbeek, K., Vanderwel, M., Vanselow, K.A., Vårhammar, A., Varone, L., Vasquez Valderrama, M.Y., Vassilev, K., Vellend, M., Veneklaas, E.J., Verbeeck, H., Verheyen, K., Vibrans, A., Vieira, I., Villacís, J., Violle, C., Vivek, P., Wagner, K., Waldram, M., Waldron, A., Walker, A.P., Waller, M., Walther, G., Wang, H., Wang, F., Wang, W., Watkins, H., Watkins, J., Weber, U., Weedon, J.T., Wei, L., Weigelt, P., Weiher, E., Wells, A.W., Wellstein, C., Wenk, E., Westoby, M., Westwood, A., White, P.J., Whitten, M., Williams, M., Winkler, D.E., Winter, K., Womack, C., Wright, I.J., Wright, S.J., Wright, J., Pinho, B.X., Ximenes, F., Yamada, T., Yamaji, K., Yanai, R., Yankov, N., Yguel, B., Zanini, K.J., Zanne, A.E., Zelený, D., Zhao, Y.P., Zheng, Jingming, Zheng, Ji, Ziemińska, K., Zirbel, C.R., Zizka, G., Zo-Bi, I.C., Zotz, G., Wirth, C., 2020. TRY plant trait database – enhanced coverage and open access. Glob. Chang. Biol. 26, 119–188. https://doi.org/10.1111/gcb.14904

Kidner, C., Umbreen, S., 2010. Why is leaf shape so variable? Int. J. Plant Dev. Biol. 4, 64–75.

Klein, L., Svoboda, H., 2017. Comprehensive Methods for Leaf Geometric Morphometric Analyses. Bio-Protocol 7, 1–21. https://doi.org/10.21769/bioprotoc.2269

Klein, L.L., Caito, M., Chapnick, C., Kitchen, C., O’Hanlon, R., Chitwood, D.H., Miller, A.J., 2017. Digital morphometrics of two north american grapevines (Vitis: Vitaceae) quantifies leaf variation between species, within species, and among individuals. Front. Plant Sci. 8, 1–10. https://doi.org/10.3389/fpls.2017.00373

Klingenberg, C.P., 2011. MorphoJ: An integrated software package for geometric morphometrics. Mol. Ecol. Resour. 11, 353–357. https://doi.org/10.1111/j.1755-0998.2010.02924.x

Kuhl, F.P., Giardina, C.R., 1982. Elliptic Fourier features of a closed contour. Comput. Graph. Image Process. 18, 236–258. https://doi.org/10.1016/0146-664X(82)90034-X

Lexer, C., Joseph, J., Van Loo, M., Prenner, G., Heinze, B., Chase, M.W., Kirkup, D., 2009. The use of digital image-based morphometrics to study the phenotypic mosaic in taxa with porous genomes. Taxon 58, 349–364. https://doi.org/10.1002/tax.582003

Li, M., An, H., Angelovici, R., Bagaza, C., Batushansky, A., Clark, L., Coneva, V., Donoghue, M.J., Edwards, E., Fajardo, D., Fang, H., Frank, M.H., Gallaher, T., Gebken, S., Hill, T., Jansky, S., Kaur, B., Klahs, P.C., Klein, L.L., Kuraparthy, V., Londo, J., Migicovsky, Z., Miller, A., Mohn, R., Myles, S., Otoni, W.C., Pires, J.C., Rieffer, E., Schmerler, S., Spriggs, E., Topp, C.N., Van Deynze, A., Zhang, K., Zhu, L., Zink, B.M., Chitwood, D.H., 2018. Topological data analysis as a morphometric method: Using persistent homology to demarcate a leaf morphospace. Front. Plant Sci. 9, 1–14. https://doi.org/10.3389/fpls.2018.00553

Lorieul, T., Pearson, K.D., Ellwood, E.R., Goëau, H., Molino, J.F., Sweeney, P.W., Yost, J.M., Sachs, J., Mata-Montero, E., Nelson, G., Soltis, P.S., Bonnet, P., Joly, A., 2019. Toward a large-scale and deep phenological stage annotation of herbarium specimens: Case studies from temperate, tropical, and equatorial floras. Appl. Plant Sci. 7. https://doi.org/10.1002/aps3.1233

Luca, G., Annamaria, G., 2019. Effectiveness of modern leaf analysis tools for the morpho-ecological study of plants: the case of *Primula albenensis*. Nord. J. Bot. 37, njb.02386. https://doi.org/10.1111/njb.02386

Manacorda, C.A., Asurmendi, S., 2018. Arabidopsis phenotyping through geometric morphometrics. Gigascience 7. https://doi.org/10.1093/gigascience/giy073

Migicovsky, Z., Li, M., Chitwood, D.H., Myles, S., 2018. Morphometrics reveals complex and heritable apple leaf shapes. Front. Plant Sci. 8, 1–14. https://doi.org/10.3389/fpls.2017.02185

Misra, S., Srivastava, A.K., Verma, S., Pandey, S., Bargali, S.S., Rana, T.S., Nair, K.N., 2017. Phenetic and genetic diversity in Indian Luffa (Cucurbitaceae) inferred from morphometric, ISSR and DAMD markers. Genet. Resour. Crop Evol. 64, 995–1010. https://doi.org/10.1007/s10722-016-0420-1

Morello, S., Sassone, A.B., López, A., 2018. Leaflet shape in the endemic South American Oxalis sect. Alpinae: An integrative approach using molecular phylogenetics and geometric morphometrics. Perspect. Plant Ecol. Evol. Syst. 35, 22–30. https://doi.org/10.1016/j.ppees.2018.09.003

Morello, S., Sede, S.M., 2016. Genetic admixture and lineage separation in a southern Andean plant. AoB Plants 8.

Nicotra, A.B., Leigh, A., Boyce, C.K., Jones, C.S., Niklas, K.J., Royer, D.L., Tsukaya, H., 2011. The evolution and functional significance of leaf shape in the angiosperms. Funct. Plant Biol. https://doi.org/10.1071/FP11057

Ostria-Gallardo, E., Ranjan, A., Chitwood, D.H., Kumar, R., Townsley, B.T., Ichihashi, Y., Corcuera, L.J., Sinha, N.R., 2016. Transcriptomic analysis suggests a key role for SQUAMOSA PROMOTER BINDING PROTEIN LIKE, NAC and YUCCA genes in the heteroblastic development of the temperate rainforest tree Gevuina avellana (Proteaceae). New Phytol. 210, 694–708. https://doi.org/10.1111/nph.13776

Page, L.M., Macfadden, B.J., Fortes, J.A., Soltis, P.S., Riccardi, G., 2015. Digitization of Biodiversity Collections Reveals Biggest Data on Biodiversity. Bioscience 65, 841–842. https://doi.org/10.1093/biosci/biv104

Pérez-Miranda, F., Mejía, O., González-Díaz, A.A., Martínez-Méndez, N., Soto-Galera, E., Zúñiga, G., Říčan, O., 2020. The role of head shape and trophic variation in the diversification of the genus Herichthys in sympatry and allopatry. J. Fish Biol. 96, 1370–1378. https://doi.org/10.1111/jfb.14304

Punyasena, S.W., Smith, S.Y., 2014. Bioinformatic and Biometric Methods in Plant Morphology. Appl. Plant Sci. 2, 1400071. https://doi.org/10.3732/apps.1400071

Renner, S., Pandey, A., 2013. The Cucurbitaceae of India: Accepted names, synonyms, geographic distribution, and information on images and DNA sequences. PhytoKeys 20, 53–118. https://doi.org/10.3897/phytokeys.20.3948

Rohlf, F.J., 2015. The tps series of software. Hystrix 26, 1–4. https://doi.org/10.4404/hystrix-26.1-11264

Royer, D.L.D., Meyerson, L.A., Robertson, K.M.K., Adams, J.M.J., 2009. Phenotypic plasticity of leaf shape along a temperature gradient in Acer rubrum. PLoS One 4, e7653. https://doi.org/10.1371/journal.pone.0007653

Schaefer, H., Renner, S.S., 2011. Phylogenetic relationships in the order Cucurbitales and a new classification of the gourd family (Cucurbitaceae). Taxon 60, 122–138. https://doi.org/10.1002/tax.601011

Schlautman, B., Diaz-Garcia, L., Barriball, S., 2020. Morphometric approaches to promote the use of exotic germplasm for improved food security and resilience to climate change: a kura clover example. Plant Sci. https://doi.org/10.1016/j.plantsci.2019.110319

Soltis, P.S., 2017. Digitization of herbaria enables novel research. Am. J. Bot. 104, 1281–1284. https://doi.org/10.3732/ajb.1700281

Soltis, P.S., Nelson, G., James, S.A., 2018. Green digitization: Online botanical collections data answering real-world questions: Online. Appl. Plant Sci. 6, 4–7. https://doi.org/10.1002/aps3.1028

Terhune, C.E., Sylvester, A.D., Scott, J.E., Ravosa, M.J., 2020. Internal architecture of the mandibular condyle of rabbits is related to dietary resistance during growth. J. Exp. Biol. 223. https://doi.org/10.1242/jeb.220988

Viscosi, V., Fortini, P., Slice, D.E., Loy, A., Blasi, C., 2009a. Geometric morphometric analyses of leaf variation in four oak species of the subgenus *Quercus* (Fagaceae). Plant Biosyst. - An Int. J. Deal. with all Asp. Plant Biol. 143, 575–587. https://doi.org/10.1080/11263500902775277

Viscosi, V., Lepais, O., Gerber, S., Fortini, P., 2009b. Leaf morphological analyses in four European oak species (Quercus) and their hybrids: A comparison of traditional and geometric morphometric methods. Plant Biosyst. 143, 564–574. https://doi.org/10.1080/11263500902723129

Wickham, H., 2016. ggplot2: Elegant Graphics for Data Analysis., ggplot2. Springer-Verlag New York. https://doi.org/10.1007/978-0-387-98141-3

Willis, C.G., Ellwood, E.R., Primack, R.B., Davis, C.C., Pearson, K.D., Gallinat, A.S., Yost, J.M., Nelson, G., Mazer, S.J., Rossington, N.L., Sparks, T.H., Soltis, P.S., 2017. Old Plants, New Tricks: Phenological Research Using Herbarium Specimens. Trends Ecol. Evol. 32, 531–546. https://doi.org/10.1016/j.tree.2017.03.015

Zhang, L.B., Simmons, M.P., Kocyan, A., Renner, S.S., 2006. Phylogeny of the Cucurbitales based on DNA sequences of nine loci from three genomes: Implications for morphological and sexual system evolution. Mol. Phylogenet. Evol. 39, 305–322. https://doi.org/10.1016/j.ympev.2005.10.002

